# Genome-Bench: A Scientific Reasoning Benchmark from Real-World Expert Discussions

**DOI:** 10.1101/2025.06.02.657538

**Authors:** Ming Yin, Yuanhao Qu, Dyllan Liu, Ling Yang, Le Cong, Mengdi Wang

## Abstract

In this short report, we present an automated pipeline tailored for the genomics domain and introduce *Genome-Bench*, a new benchmark constructed from over a decade of scientific forum discussions on genome engineering. Our pipeline transforms raw interactions into a reinforcement learningfriendly multiple-choice questions format, supported by 3000+ high-quality question-answer pairs spanning foundational biology, experimental troubleshooting, tool usage, and beyond. To our knowledge, this is the first end-to-end pipeline for teaching LLMs to reason from scientific discussions, with promising potential for generalization across scientific domains beyond biology. The dataset is available at https://huggingface.co/datasets/Mingyin0312/Genome-Bench.

## 1 Introduction

Large language models (LLMs) have shown remarkable capabilities in domains such as mathematics [7, 13] and programming [2]. However, despite their impressive performance on standardized reasoning benchmarks, LLMs still struggle with expert-level scientific reasoningparticularly in complex, high-stakes fields such as genomics. Unlike mathematical proofs or coding problems, which typically have clear-cut solutions and deterministic evaluation criteria, scientific reasoning often requires interpreting ambiguous observations, accounting for domain-specific nuances, and synthesizing knowledge across multiple layers of abstraction. This gap between current LLM capabilities and the level of reasoning expected from domain experts presents a major challenge for applying LLMs to real-world scientific workflows.

To bridge the gap between LLMs and human expertise, we demonstrate an end-to-end pipeline for training LLMs based on *authentic* scientific discussions among human experts in the field of genomics. We introduce **Genome-Bench**, a scientific reasoning benchmark in genomics comprising of questions and answers extracted from over a decade of archived scientific forum discussions among genome editing practitioners. These open discussions, drawn from a Google Research forum established in 2013 to explore CRISPR gene-editing technologies, reflect the nuanced, real-world problem-solving and methodological decision-making that working scientists engage in. Unlike curated benchmarks like GPQA [12], HLE [11], and Lab-Bench [6], which often emphasize standardized testing for LLMs, Genome-Bench captures authentic scientific reasoning as it unfolds in practice. It offers a unique lens into how scientists navigate common misconceptions, interpret ambiguous results, and refine experimental protocols over time. Our main contributions are summarized as follows:

Technical Report.

- **Genome-Bench: A novel benchmark grounded in real expert discussions**. We present the first LLM benchmark constructed from real-world scientific discussions, capturing the reasoning processes of practicing researchers. Genome-Bench comprises 3,332 meticulously curated multiple-choice questions derived from over a decade of CRISPR forum archives. Unlike synthetic or exam-based datasets, Genome-Bench reflects the genuine complexity and problem-solving strategies found in frontline experimental biology. The questions span diverse subfieldsfrom experimental troubleshooting and reagent selection to protocol design and tool interpretationproviding a rich and realistic testbed for evaluating domain-specific reasoning in LLMs.
- **An end-to-end data pipeline for transforming expert dialogue into structured RL-compatible data**. We develop an automated end-to-end pipeline that converts raw, noisy discussion email threads into structured question-answer pairs suitable for training and evaluating LLMs with reinforcement learning. This pipeline integrates LLM-assisted question-answer extraction, distractor generation, context preservation, and rigorous quality control to produce high-quality multiple-choice questions and rule-based rewards with expert-style rationales. This framework is modular and domain-agnostic, enabling scalable construction of reasoning datasets across scientific domains.

## 2 Related Works

In biomedical and scientific domains, a variety of QA datasets have been curated to facilitate the development and evaluation of domain-specialized models. The BioASQ challenge [5] provides a benchmark for biomedical question answering with expert-annotated answers and linked snippets from literature [14]. In the medical domain, PubMedQA [4] targets fact-based questions over biomedical abstracts, MedMCQA [9] spans clinical specialties, and MedQA [3] and COVID-QA

[8] address disease diagnosis and pandemic-related questions, respectively. emrQA [10] focuses on EMR-based question answering, and MASH-QA [15] explores long, multi-span answers. These datasets typically derive from formal sources such as exams, medical records, or curated biomedical corpora. More recently, LAB-Bench [6] has been introduced as a comprehensive benchmark to evaluates LLMs on real-world biology research taskssuch as literature search, protocol planning, and data analysisoffering a more practical alternative to textbook-style QA datasets. In contrast, our work introduces the first QA/MCQ dataset centered on CRISPR gene editing and constructed from authentic scientific discussions.

## 3 Genome-Bench

We introduce Genome-Bench dataset. An overview of the full data pipeline is shown in Figure 1.

**Figure 1.**
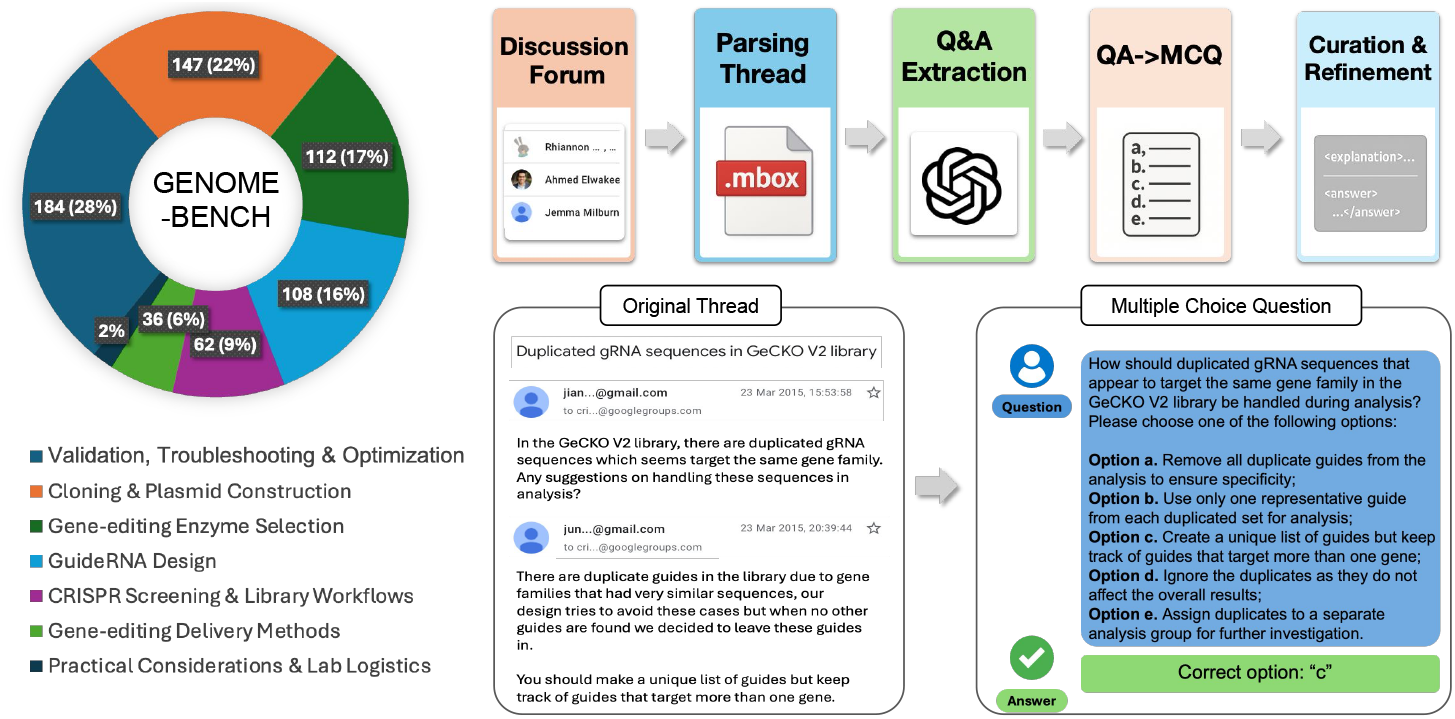
*Overview of Genome-Bench*. We introduce GENOME-BENCH, a novel scientific bench-mark comprising 3,332 question-answer pairs derived directly from real-world genome research discussions among human experts over 11 years. The dataset uniquely captures authentic scientific discourse spanning seven key areasfrom experimental Validation and Troubleshooting to Practical Considerations and Lab Logistics. Our end-to-end data pipeline begins with parsing raw email threads from archived scientific forums in .mbox format, leveraging a Large Language Model (LLM) to systematically extract question-answer-context tuples. These tuples are then transformed into a structured multiple-choice format, meticulously curated through deduplication, quality filtering, and precise answer annotation to ensure high-quality benchmarking for genomic research applications.

**Figure 2.**
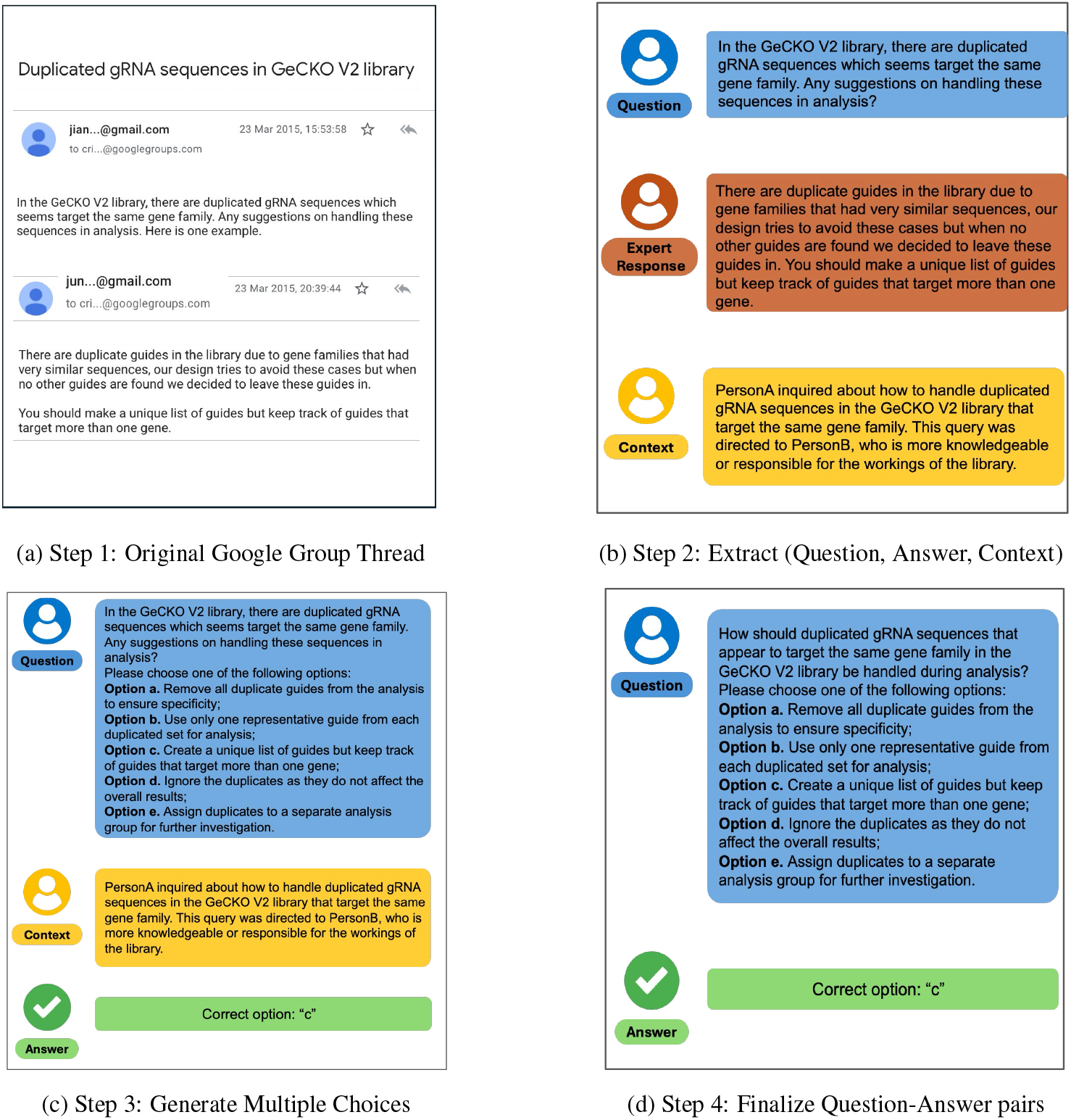
A Step-by-step Example of Genome-Bench Creation. This figure illustrates the data extraction pipeline for curating the Genome-Bench. Initially, the original discussion thread is parsed into a structured format consisting of a Question, Expert Response, and Context, where the context is carefully crafted to reduce potential hallucinations. The expert’s response is subsequently transformed into a multiple-choice question, with the correct answer directly derived from the expert’s original reply. Finally, the QA pair is refined by integrating the context into the initial question.

### Data Source and Significance

Genome-Bench is a new benchmark dataset designed to evaluate and advance scientific reasoning in large language models (LLMs), constructed from over a decade of CRISPR-related genome discussions [1]. The source material originates from a long-standing public mailing list established by researchers at the Broad Institute of Harvard and MIT. Spanning 20132023, this forum captured thousands of real-world scientific inquiries and peer responses related to gene editing. Unlike curated educational datasets or exam-derived corpora, Genome-Bench reflects how scientists actually reason in the lab: asking open-ended, ambiguous questions, proposing competing hypotheses, and resolving experimental uncertainties collaboratively. The resulting dataset comprises of high-quality questionanswer pairs, touching on diverse subfields including foundational biology, experimental troubleshooting, protocol optimization, reagent selection, tool usage, and lab logistics. Because the content is grounded in expert-to-expert conversations, it captures nuanced reasoning patterns, contextual dependencies, and even common misconceptions that synthetic QA datasets often miss.

### An End-to-End Data Processing Pipeline

To transform this corpus into a benchmark suitable for LLM training and evaluation, we developed a fully automated pipeline that processes raw .mbox email archives into structured *multiple-choice questions*. The pipeline includes the following stages:

1. *Thread Parsing and Q&A Extraction*. Each email thread is parsed and preprocessed using a custom parser and GPT-4-Turbo. For each thread, the model extracts a concise scientific question, an expert-provided answer, and supporting context from the surrounding discussion. The result is a structured triplet: (question, answer, context). This ensures that questions retain their real-world framing such as lab-specific constraints and prior experimental attempts.
2. *MCQ Generation*. The extracted tuples are converted into single-answer MCQs. First, contextual information is prepended to the question to ensure self-containment. Then, the question is rewritten into a fluent, standalone prompt using few-shot GPT-4o instructions. Plausible distractors are generated using LLM prompting, guided to produce scientifically credible but incorrect alternatives that mirror realistic errors. All answer choices are randomly shuffled, with the correct answer explicitly labeled for evaluation or training.
3. *Answer Encoding and Formatting*. Each item is serialized into a standardized format with consistent labeling. Correct answers are marked (ae), and expert reasoning is enclosed in <explanation>…</explanation> tags, followed by a final answer tag (<answer>…</answer>). This format facilitates both supervised fine-tuning and reinforcement learning, as it provides both the correct answer and the rationale behind it.
4. *Quality Filtering*. We applied a multi-stage quality control process. Duplicate or near-duplicate entries were removed, as were questions with ambiguous or trivial answers. Additional filtering excluded off-topic posts and threads lacking substantive expert engagement. The final dataset consists of 3,332 verified, high-quality QA items.

### Benchmark Structure

The final 3,332 questions was partitioned into **a training set** and **a test set**, comprising 2,671 and 661 questions, respectively. To enable detailed evaluation of LLM scientific reasoning, each Genome-Bench test question is annotated with two additional fields: *category* and *difficulty*. We defined seven thematic categories based on a corpus-wide analysis: *Validation, Troubleshooting & Optimization, Cloning & Plasmid Construction, Gene-editing Enzyme Selection, GuideRNA Design, Screening & Library Design, Gene-editing Delivery Methods* and *Practical Lab Logistics*. These categories reflect recurring topics in experimental CRISPR work and enable domain-specific performance breakdowns. In parallel, questions are labeled with difficulty levels*Easy, Medium, or Hard*based on linguistic structure and conceptual complexity. Heuristics used for this annotation include the presence of conditional clauses, the depth of reasoning required, and the presence of multiple plausible distractors. This stratification enables fine-grained evaluation of model robustness as cognitive load increases. We use training data for fine-tuning and test data for evaluation in the following sections.

## 4 Conclusion

In this paper, we present Genome-Bench, a novel benchmark for scientific reasoning in genomics, built from over a decade of authentic CRISPR forum discussions and comprising 3,300+ expertly curated multiple-choice questions. Our end-to-end pipeline transforms raw expert dialogue into structured QA data suitable for reinforcement learning, capturing the real-world complexity, ambiguity, and nuance of experimental biologyunlike synthetic or exam-based datasets. By grounding evaluation in authentic scientific discourse, Genome-Bench provides a realistic testbed for domain-specific performance. Its modular and domain-agnostic design enables extension to other scientific fields, offering a scalable framework for constructing practical, reasoning-oriented benchmarks that bring LLMs closer to integration in real-world laboratory workflows.

## Supporting information

Source Latex Code and Figures

## Appendix

### A Detailed pipeline for Constructing the Genome-Bench Dataset

#### Data Source

The source of our dataset is an open, public discussion forum “Genome Engineering using CRISPR/Cas Systems,” [1] initially established by the Feng Zhang lab at the Broad Institute of MIT and Harvard. This forum served as a dynamic, crowd-sourced Q&A platform where scientists worldwide could post questions about CRISPR gene-editing tools and laboratory practices. Over 11 years, it amassed a wealth of inquiries and expert responses, culminating in approximately 4,000 discussion threads.

#### Data Parsing and Q&A Extraction

The raw dataset, exported in `.mbox`, is parsed and converted into DataFrame format using Pandas, where each row corresponds to an email thread. Each unique email thread is individually pre-processed by OpenAIs GPT-4 Turbo model and reformatted for the purpose of fine-tuning. The model is tasked with extracting Q&A pairs by interpreting the textual content of each thread. The model is prompted to process the current email thread and identify scientific and research related questions and answers (Q&A Pairs). Because certain Q&As are specific and context-driven, the model is prompted to use the entire thread to provide a “context” field for each Q&A Pair. It finally outputs structured data with the following format: `{question, answer, context}`. The Q&A dataset is anonymized after processing.

#### Q&A Format to Multiple-Choice Question (MCQ) Transformation

To utilize the data for training and evaluation, we converted each Q&A pair into a structured multiple-choice question format. The transformation was carefully designed to preserve the original problem and answer while adding plausible alternative options to form a challenging MCQ. Our pipeline consists of several sequential steps:

##### Context Integration

Many forum questions rely on context from previous messages (e.g., experimental conditions, what has been tried already). We prepend any available context to the question to make it self-contained. For example, if a question was asked in a thread after a description of an experiment, we combine them: `“Question context: … [context description] … Question: … [actual question]”`. This ensures that important background information is not lost.

##### Question Rewriting

We then convert the question (with context) into a natural, stand-alone question that reads clearly on its own. This involves rewriting to integrate the context smoothly and remove any forum-specific phrasing (such as direct references to users like “Person A”). We employed a strong LLM (GPT-4o in our case) to perform this rewriting via prompt-based instruction. The prompt asked the model to preserve all factual information while phrasing the question in a single, coherent sentence or paragraph. For instance, the structured input `“Question context: … PersonB suggests … PersonA addresses. Question: Is it` `possible your Amp/Carbenicillin has gone bad?”` was rewritten by GPT-4 to: `Could the issue with your lentiCrispr v2 growth after cloning the gRNA be due to your ampicillin or carbenicillin going bad?` a fluent question that incorporates the context. This step yields a polished question ready for future steps.

##### Distractor Generation

Next, we generate four distractor options (incorrect answers) to accompany the correct answer. The correct answer is derived from the original answer text or its key conclusion. We prompt GPT-4o with the question and the correct answer (explanation) to produce plausible alternative answers that could be chosen by a non-expert or a confused model. The model is instructed to produce options that are credible yet definitively incorrect given the context. For example, if the correct answer explains that a plasmids large size likely caused a transfection failure, distractors might include other causes like reagent issues or claims that plasmid size doesnt matter. The generation is done in a prompt that asks for five candidate answers (one being the true answer, and four false ones). We then identify which of the models outputs corresponds to the known correct answer and label it accordingly. This step yields a set of five answer statements (ae).

##### Option Formatting and Shuffling

We format the answer options as lettered choices prefixed by `a., b., c., d., e.`, and ensure they follow a consistent style, and we randomize the order of the options. The correct answers label is tracked during shuffling so that we know which letter is correct after permutation. Shuffling prevents positional bias and makes the dataset more robust (see Figure 3). We merge the rewritten question and the shuffled options into one consolidated question text. In the final dataset, the question field contains both the prompt and the available choices. For example, `Question: Could the issue with your lentiCrispr v2 growth be due to your Ampicillin or Carbenicillin going bad? Please choose one of the following options: a. b. c. d. e.`. This step ensures each data entry is a complete multiple-choice question ready to be posed to a model or human.

**Figure 3.**
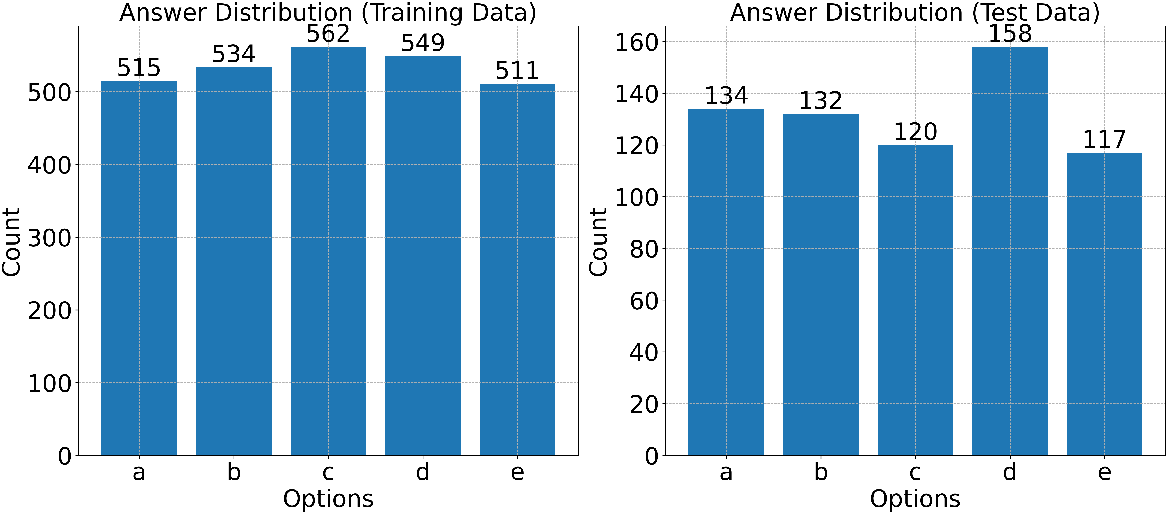
Answer distribution for Genome-Bench.

##### Answer and Explanation Encoding

We construct an answer field containing the correct option label and the explanation. We wrap the original answer explanation (usually a few sentences of reasoning from the expert) in `<explanation>…</explanation>` tags, and we place the correct option letter in `<answer>…</answer>` tags. For instance, an entrys answer might be: `<explanation>Yes, larger plasmid size can reduce transfection efficiency in certain cell lines, so using too much DNA likely caused the issue.</explanation> <answer>e</answer>`. The explanation text is taken directly from the experts answer, ensuring we preserve the reasoning behind the correct choice.

After these steps, each data sample is a JSON object with a question (including integrated context and all answer options) and an answer (containing the correct choice and explanation). The dataset is thus in a structured QA format suitable for both training an LLM (where the model is given the question and must output the answer) and evaluating its multiple-choice accuracy. An example from our training set is shown below:

##### Question

In the process of using CRISPR technology on plants, particularly when considering whether these plants can be classified as cisgenic, isn’t there a possibility that some T-DNA sequence from the plant binary vector backbone might insert into the plant genome during Agrobacterium-mediated transformation?

Please choose one of the following options:

- a. Only in rare cases, the T-DNA sequence from the binary vector may insert into the plant genome.
- b. T-DNA sequences are never inserted into the plant genome during transformation.
- c. The plant genome naturally resists any insertion of T-DNA sequences from the binary vector backbone.
- d. Yes, the T-DNA sequence from the binary vector backbone typically inserts into the plant genome.
- e. No, T-DNA insertion is highly selective and does not include the binary vector backbone.

##### Answer

<explanation>Yes, during Agrobacterium-mediated transformation, some T-DNA from the plant binary vector backbone typically inserts into the plant genome.</explanation> <answer>d</answer>

#### Quality Control

Because the raw mined MCQ can be noisy, we performed several quality control and filtering steps to clean the dataset before use:

##### Deduplication

We removed duplicate or nearly identical questions to avoid redundancy. Some questions were asked multiple times over the years; these were identified (using string matching and manual verification) and only one representative instance was kept.

##### Incomplete/Unanswered Removal

We filtered out any questions that did not have a clear answer in the discussion. Specifically, entries where the extracted answer was empty, merely a confirmation, or contained phrases indicating no answer (e.g., No answer, unanswered, no response) were discarded. In total, we removed on the order of 180+ Q&A pairs in this category, which had no informative value due to lack of resolution.

##### Low-Quality Filtering

We also pruned questions that were overly vague, off-topic, or lacking scientific substance, as identified by human inspection. These included generic pleas for help (Can someone please give suggestions what could be going wrong?), questions about website availability and collaboration requests.

After cleaning, we arrived at a finalized set of 3,332 single-choice multiple-choice questions, which we split into a training set and a test set. We reserved 20% for testing, resulting in 2,671 training questions and 661 test questions. An additional example is also visualized in Figure 2.

#### Genome-Bench Evaluation Protocol

To ensure consistency across all evaluations, we employed a structured system prompt that specifies the required response format. The model is instructed to first generate an internal reasoning process enclosed within `<explanation>…</explanation>` tags, followed by its final answer enclosed within `<answer>…</answer>` tags. The final answer must be exactly one of five choices: `{a, b, c, d, e}`. This format ensures that models not only select an answer but also provide traceable reasoning steps.

##### Answer Extraction

After inference, each generated output was parsed to extract the final answer. We implemented a simple parser that locates the text enclosed within the <answer>…</answer> tags. In cases where the expected format was not adhered to, the output was considered invalid and treated accordingly during evaluation.

