## Supplementary material for "Genome-Bench: A Scientific Reasoning Benchmark from Real-World Expert Discussions": Source Latex Code and Figures: appendix_1.pdf

### Duplicated gRNA sequences in GeCKO V2 library

****
to

23 Mar 2015, 15:53:58

In the GeCKO V2 library, there are duplicated gRNA sequences which seems target the same gene family. Any suggestions on handling these sequences in analysis. Here is one example.

****
to

23 Mar 2015, 20:39:44

There are duplicate guides in the library due to gene families that had very similar sequences, our design tries to avoid these cases but when no other guides are found we decided to leave these guides in.

You should make a unique list of guides but keep track of guides that target more than one gene.
