## Supplementary material for "Genome-Bench: A Scientific Reasoning Benchmark from Real-World Expert Discussions": Source Latex Code and Figures: appendix_1_3.pdf

**Question**

In the GeCKO V2 library, there are duplicated gRNA sequences which seems target the same gene family. Any suggestions on handling these sequences in analysis?
Please choose one of the following options:
**Option a.** Remove all duplicate guides from the analysis to ensure specificity;
**Option b.** Use only one representative guide from each duplicated set for analysis;
**Option c.** Create a unique list of guides but keep track of guides that target more than one gene;
**Option d.** Ignore the duplicates as they do not affect the overall results;
**Option e.** Assign duplicates to a separate analysis group for further investigation.

**Context**

PersonA inquired about how to handle duplicated gRNA sequences in the GeCKO V2 library that target the same gene family. This query was directed to PersonB, who is more knowledgeable or responsible for the workings of the library.

**Answer**

Correct option: “c”
