## Supplementary material for "Genome-Bench: A Scientific Reasoning Benchmark from Real-World Expert Discussions": Source Latex Code and Figures: appendix_1_4.pdf

**Question**

How should duplicated gRNA sequences that appear to target the same gene family in the GeCKO V2 library be handled during analysis? Please choose one of the following options:

**Option a.** Remove all duplicate guides from the analysis to ensure specificity;

**Option b.** Use only one representative guide from each duplicated set for analysis;

**Option c.** Create a unique list of guides but keep track of guides that target more than one gene;

**Option d.** Ignore the duplicates as they do not affect the overall results;

**Option e.** Assign duplicates to a separate analysis group for further investigation.

**Answer**

Correct option: "c"
