## Supplementary material for "Genome-Bench: A Scientific Reasoning Benchmark from Real-World Expert Discussions": Source Latex Code and Figures: human_temp1.pdf

In our analysis of the CRISPR/Cas9 gene screening results, we noticed something surprising: while one of the six sgRNAs targeting the same gene confers resistance and is enriched, another one causes the population to drop out. Does anyone know why this might be happening?

a. "All sgRNAs targeting the same gene should have identical effects"
b. "The guide RNAs are not binding to the target gene at all"
c. "The Cas9 enzyme used is defective in these experiments"
d. "The sgRNAs may have off-target effects that cause different outcomes"
e. "Gene editing always leads to consistent outcomes regardless of sgRNA used”

**Human Answer:** d
**Human Explanation:** Though targeting at the same gene, sgRNAs might have different on-target and off-target efficiency. The most common cause of population drop out is due to the genotoxicity cased by the significant off-target effects.

**Model Answer:** d
**Model Explanation:** sgRNAs are programmed to bind to specific sequences. If two sgRNAs target the same gene, but the second one has off-target effects, it could cause the population to drop out. Off-target effects occur when the sgRNA binds to a different location.

### Partially Wrong Reasoning

Comment: Model’s explanation on the off-target effect as the cause, but incorrectly describe the relation on off-target effects and dropout phenotype.
