## Supplementary material for "Genome-Bench: A Scientific Reasoning Benchmark from Real-World Expert Discussions": Source Latex Code and Figures: human_temp2.pdf

In the context of using CRISPR for tagging a protein at the DNA level, if the guide RNA we use also targets the donor plasmid, will this completely eliminate homologous recombination, or will it just reduce its frequency? Please choose one of the following options:

a. "It will completely eliminate HR"
b. "It will only cause a temporary delay in HR"
c. "It will not affect HR at all"
d. "It will increase the frequency of HR"
e. "It will reduce the frequency of HR"

**Human Answer:** e
**Human Explanation:** Guide RNA would cleave the donor plasmid which will lead to significant reduction of HR frequency. While, cleavage of all donor plasmids won't happen simultaneously so it's likely to still have some HR event.

**Model Answer:** e
**Model Explanation:** When the guide RNA used also targets the donor plasmid, it will cause the plasmid to be degraded before it can contribute to homologous recombination. This will completely prevent homologous recombination caused by the plasmid.

### Completely Wrong Reasoning

Comment: The explanation model provides refers to a different answer in option.
