## Supplementary material for "Genome-Bench: A Scientific Reasoning Benchmark from Real-World Expert Discussions": Source Latex Code and Figures: human_temp.pdf

Can you explain how I might use GFP to indicate the transfection efficiency of a Px330 CRISPR vector into a cell line, and what transduction methods might be recommended? Please choose one of the following options:

a. "By measuring the fluorescence intensity of CRISPR vector"
b. "By integrating GFP directly into the CRISPR vector"
c. "By using GFP-tagged Cas9 protein"
d. "By observing cell morphology changes under a GFP filter”
e. "By using another commercial vector that carries GFP as a transfection control and transfecting it in another well concurrently with transfecting your CRISPR vector"

**Human Answer:** e
**Human Explanation:** Doing the co-transfection with a GFP vector could be an easy way to measure the transfection efficiency of the CRISPR vector without modifying the CRISPR vector.

**Model Answer:** e
**Model Explanation:** When it comes to assessing the transfection efficiency of a CRISPR vector, it's best to use a separate vector like GFP as a transfection control, and transfect them both at the same time to optimize transfection conditions.

### Correct Reasoning

Comment: Model selects the right answer and provided correct reasoning.
