## Supplementary material for "Genome-Bench: A Scientific Reasoning Benchmark from Real-World Expert Discussions": Source Latex Code and Figures: ref.bib

@article{discussion,
  title={Public Google Discussion Group: Genome Engineering CRISPR/Cas Systems},
  note={\url{https://groups.google.com/g/crispr?pli=1}}
}

@article{shao2024deepseekmath,
  title={Deepseekmath: Pushing the limits of mathematical reasoning in open language models},
  author={Shao, Zhihong and Wang, Peiyi and Zhu, Qihao and Xu, Runxin and Song, Junxiao and Bi, Xiao and Zhang, Haowei and Zhang, Mingchuan and Li, YK and Wu, Y and others},
  journal={arXiv preprint arXiv:2402.03300},
  year={2024}
}

@article{jiang2024survey,
  title={A survey on large language models for code generation},
  author={Jiang, Juyong and Wang, Fan and Shen, Jiasi and Kim, Sungju and Kim, Sunghun},
  journal={arXiv preprint arXiv:2406.00515},
  year={2024}
}

@article{schulman2017proximal,
  title={Proximal policy optimization algorithms},
  author={Schulman, John and Wolski, Filip and Dhariwal, Prafulla and Radford, Alec and Klimov, Oleg},
  journal={arXiv preprint arXiv:1707.06347},
  year={2017}
}

@article{krithara2023bioasq,
  title={BioASQ-QA: A manually curated corpus for Biomedical Question Answering},
  author={Krithara, Anastasia and Nentidis, Anastasios and Bougiatiotis, Konstantinos and Paliouras, Georgios},
  journal={Scientific Data},
  volume={10},
  number={1},
  pages={170},
  year={2023},
  publisher={Nature Publishing Group UK London}
}

@article{zhang2025scientific,
  title={Scientific large language models: A survey on biological \& chemical domains},
  author={Zhang, Qiang and Ding, Keyan and Lv, Tianwen and Wang, Xinda and Yin, Qingyu and Zhang, Yiwen and Yu, Jing and Wang, Yuhao and Li, Xiaotong and Xiang, Zhuoyi and others},
  journal={ACM Computing Surveys},
  volume={57},
  number={6},
  pages={1--38},
  year={2025},
  publisher={ACM New York, NY}
}

@article{prabhakar2025omniscience,
  title={OmniScience: A Domain-Specialized LLM for Scientific Reasoning and Discovery},
  author={Prabhakar, Vignesh and Islam, Md Amirul and Atanas, Adam and Wang, Yao-Ting and Han, Joah and Jhunjhunwala, Aastha and Apte, Rucha and Clark, Robert and Xu, Kang and Wang, Zihan and others},
  journal={arXiv preprint arXiv:2503.17604},
  year={2025}
}

@article{liang2024mapping,
  title={Mapping the increasing use of LLMs in scientific papers},
  author={Liang, Weixin and Zhang, Yaohui and Wu, Zhengxuan and Lepp, Haley and Ji, Wenlong and Zhao, Xuandong and Cao, Hancheng and Liu, Sheng and He, Siyu and Huang, Zhi and others},
  journal={arXiv preprint arXiv:2404.01268},
  year={2024}
}

@article{xie2023darwin,
  title={Darwin series: Domain specific large language models for natural science},
  author={Xie, Tong and Wan, Yuwei and Huang, Wei and Yin, Zhenyu and Liu, Yixuan and Wang, Shaozhou and Linghu, Qingyuan and Kit, Chunyu and Grazian, Clara and Zhang, Wenjie and others},
  journal={arXiv preprint arXiv:2308.13565},
  year={2023}
}

@article{pal2024domain,
  title={A domain-specific next-generation large language model (LLM) or ChatGPT is required for biomedical engineering and research},
  author={Pal, Soumen and Bhattacharya, Manojit and Lee, Sang-Soo and Chakraborty, Chiranjib},
  journal={Annals of biomedical engineering},
  volume={52},
  number={3},
  pages={451--454},
  year={2024},
  publisher={Springer}
}

@article{cai2024sciassess,
  title={Sciassess: Benchmarking llm proficiency in scientific literature analysis},
  author={Cai, Hengxing and Cai, Xiaochen and Chang, Junhan and Li, Sihang and Yao, Lin and Wang, Changxin and Gao, Zhifeng and Wang, Hongshuai and Li, Yongge and Lin, Mujie and others},
  journal={arXiv preprint arXiv:2403.01976},
  year={2024}
}

@article{luo2023biomedgpt,
  title={Biomedgpt: Open multimodal generative pre-trained transformer for biomedicine},
  author={Luo, Yizhen and Zhang, Jiahuan and Fan, Siqi and Yang, Kai and Wu, Yushuai and Qiao, Mu and Nie, Zaiqing},
  journal={arXiv preprint arXiv:2308.09442},
  year={2023}
}

@article{bai2022training,
  title={Training a helpful and harmless assistant with reinforcement learning from human feedback},
  author={Bai, Yuntao and Jones, Andy and Ndousse, Kamal and Askell, Amanda and Chen, Anna and DasSarma, Nova and Drain, Dawn and Fort, Stanislav and Ganguli, Deep and Henighan, Tom and others},
  journal={arXiv preprint arXiv:2204.05862},
  year={2022}
}

@article{zhang2024generalist,
  title={A generalist vision--language foundation model for diverse biomedical tasks},
  author={Zhang, Kai and Zhou, Rong and Adhikarla, Eashan and Yan, Zhiling and Liu, Yixin and Yu, Jun and Liu, Zhengliang and Chen, Xun and Davison, Brian D and Ren, Hui and others},
  journal={Nature Medicine},
  pages={1--13},
  year={2024},
  publisher={Nature Publishing Group US New York}
}

@article{guo2025deepseekr1,
  title={Deepseek-r1: Incentivizing reasoning capability in llms via reinforcement learning},
  author={Guo, Daya and Yang, Dejian and Zhang, Haowei and Song, Junxiao and Zhang, Ruoyu and Xu, Runxin and Zhu, Qihao and Ma, Shirong and Wang, Peiyi and Bi, Xiao and others},
  journal={arXiv preprint arXiv:2501.12948},
  year={2025}
}

@article{kumar2024training,
	title={Training language models to self-correct via reinforcement learning},
	author={Kumar, Aviral and Zhuang, Vincent and Agarwal, Rishabh and Su, Yi and Co-Reyes, John D and Singh, Avi and Baumli, Kate and Iqbal, Shariq and Bishop, Colton and Roelofs, Rebecca and others},
	journal={arXiv preprint arXiv:2409.12917},
	year={2024}
}

@article{shinn2024reflexion,
	title={Reflexion: Language agents with verbal reinforcement learning},
	author={Shinn, Noah and Cassano, Federico and Gopinath, Ashwin and Narasimhan, Karthik and Yao, Shunyu},
	journal={Advances in Neural Information Processing Systems},
	volume={36},
	year={2024}
}

@article{yang2025reasonflux,
  title={ReasonFlux: Hierarchical LLM Reasoning via Scaling Thought Templates},
  author={Yang, Ling and Yu, Zhaochen and Cui, Bin and Wang, Mengdi},
  journal={arXiv preprint arXiv:2502.06772},
  year={2025}
}

@inproceedings{wang2024math,
  title={Math-shepherd: Verify and reinforce llms step-by-step without human annotations},
  author={Wang, Peiyi and Li, Lei and Shao, Zhihong and Xu, Runxin and Dai, Damai and Li, Yifei and Chen, Deli and Wu, Yu and Sui, Zhifang},
  booktitle={Proceedings of the 62nd Annual Meeting of the Association for Computational Linguistics (Volume 1: Long Papers)},
  pages={9426--9439},
  year={2024}
}

@article{r1-searcher,
  title={R1-Searcher: Incentivizing the Search Capability in LLMs via Reinforcement Learning},
  author={Song, Huatong and Jiang, Jinhao and Min, Yingqian and Chen, Jie and Chen, Zhipeng and Zhao, Wayne Xin and Fang, Lei and Wen, Ji-Rong},
  journal={arXiv preprint arXiv:2503.05592},
  year={2025}
}

@article{yu2025dapo,
  title={Dapo: An open-source llm reinforcement learning system at scale},
  author={Yu, Qiying and Zhang, Zheng and Zhu, Ruofei and Yuan, Yufeng and Zuo, Xiaochen and Yue, Yu and Fan, Tiantian and Liu, Gaohong and Liu, Lingjun and Liu, Xin and others},
  journal={arXiv preprint arXiv:2503.14476},
  year={2025}
}

@article{wu2023pairwise,
  title={Pairwise proximal policy optimization: Harnessing relative feedback for llm alignment},
  author={Wu, Tianhao and Zhu, Banghua and Zhang, Ruoyu and Wen, Zhaojin and Ramchandran, Kannan and Jiao, Jiantao},
  journal={arXiv preprint arXiv:2310.00212},
  year={2023}
}

@article{zhong2024dpo,
  title={Dpo meets ppo: Reinforced token optimization for rlhf},
  author={Zhong, Han and Shan, Zikang and Feng, Guhao and Xiong, Wei and Cheng, Xinle and Zhao, Li and He, Di and Bian, Jiang and Wang, Liwei},
  journal={arXiv preprint arXiv:2404.18922},
  year={2024}
}

@article{chang2023learning,
  title={Learning to generate better than your llm},
  author={Chang, Jonathan D and Brantley, Kiante and Ramamurthy, Rajkumar and Misra, Dipendra and Sun, Wen},
  journal={arXiv preprint arXiv:2306.11816},
  year={2023}
}


@article{tsatsaronis2015overview,
  title={An overview of the BIOASQ large-scale biomedical semantic indexing and question answering competition},
  author={Tsatsaronis, George and Balikas, Georgios and Malakasiotis, Prodromos and Partalas, Ioannis and Zschunke, Matthias and Alvers, Michael R and Weissenborn, Dirk and Krithara, Anastasia and Petridis, Sergios and Polychronopoulos, Dimitris and others},
  journal={BMC bioinformatics},
  volume={16},
  pages={1--28},
  year={2015},
  publisher={Springer}
}

@article{jin2019pubmedqa,
  title={Pubmedqa: A dataset for biomedical research question answering},
  author={Jin, Qiao and Dhingra, Bhuwan and Liu, Zhengping and Cohen, William W and Lu, Xinghua},
  journal={arXiv preprint arXiv:1909.06146},
  year={2019}
}

@inproceedings{pal2022medmcqa,
  title={Medmcqa: A large-scale multi-subject multi-choice dataset for medical domain question answering},
  author={Pal, Ankit and Umapathi, Logesh Kumar and Sankarasubbu, Malaikannan},
  booktitle={Conference on health, inference, and learning},
  pages={248--260},
  year={2022},
  organization={PMLR}
}

@article{jin2021disease,
  title={What disease does this patient have? a large-scale open domain question answering dataset from medical exams},
  author={Jin, Di and Pan, Eileen and Oufattole, Nassim and Weng, Wei-Hung and Fang, Hanyi and Szolovits, Peter},
  journal={Applied Sciences},
  volume={11},
  number={14},
  pages={6421},
  year={2021},
  publisher={MDPI}
}

@article{yang2022gatortron,
  title={GatorTron: A Large Clinical Language Model to Unlock Patient Information from Unstructured Electronic Health Records},
  author={Yang, Xi and Chen, Aokun and PourNejatian, Nima and Shin, Hoo Chang and Smith, Kaleb E. and Parisien, Christopher and Compas, Colin and Martin, Cheryl and Flores, Mona G. and Zhang, Ying and Magoc, Tanja and Harle, Christopher A. and Lipori, Gloria and Mitchell, Duane A. and Hogan, William R. and Shenkman, Elizabeth A. and Bian, Jiang and Wu, Yonghui},
  journal={arXiv preprint arXiv:2203.03540},
  year={2022},
  url={https://arxiv.org/abs/2203.03540}
}

@inproceedings{moller2020covidqa,
  title={COVID-QA: A question answering dataset for COVID-19},
  author={M{\"o}ller, Timo and Reina, Anthony and Jayakumar, Raghavan and Pietsch, Malte},
  booktitle={Proceedings of the 1st Workshop on NLP for COVID-19 at ACL 2020},
  year={2020}
}

@article{pampari2018emrqa,
  title={emrqa: A large corpus for question answering on electronic medical records},
  author={Pampari, Anusri and Raghavan, Preethi and Liang, Jennifer and Peng, Jian},
  journal={arXiv preprint arXiv:1809.00732},
  year={2018}
}

@inproceedings{zhu2020question,
  title={Question answering with long multiple-span answers},
  author={Zhu, Ming and Ahuja, Aman and Juan, Da-Cheng and Wei, Wei and Reddy, Chandan K},
  booktitle={Findings of the Association for Computational Linguistics: EMNLP 2020},
  pages={3840--3849},
  year={2020}
}

@article{lee2020biobert,
  title={BioBERT: a pre-trained biomedical language representation model for biomedical text mining},
  author={Lee, Jinhyuk and Yoon, Wonjin and Kim, Sungdong and Kim, Donghyeon and Kim, Sunkyu and So, Chan Ho and Kang, Jaewoo},
  journal={Bioinformatics},
  volume={36},
  number={4},
  pages={1234--1240},
  year={2020},
  publisher={Oxford University Press}
}

@article{beltagy2019scibert,
  title={SciBERT: A pretrained language model for scientific text},
  author={Beltagy, Iz and Lo, Kyle and Cohan, Arman},
  journal={arXiv preprint arXiv:1903.10676},
  year={2019}
}

@article{taylor2022galactica,
  title={Galactica: A large language model for science},
  author={Taylor, Ross and Kardas, Marcin and Cucurull, Guillem and Scialom, Thomas and Hartshorn, Anthony and Saravia, Elvis and Poulton, Andrew and Kerkez, Viktor and Stojnic, Robert},
  journal={arXiv preprint arXiv:2211.09085},
  year={2022}
}

@article{huang2024crispr,
  title={Crispr-gpt: An llm agent for automated design of gene-editing experiments},
  author={Huang, Kaixuan and Qu, Yuanhao and Yin, Ming and Zhan, Kanghong and Liu, Dyllan and Yin, Di and Johnson, William A and  Zhou, Denny and Altman, Russ and Wang, Mengdi and Cong, Le},
  journal={nature Biomedical Engineering},
  year={2025}
}

@article{luo2022biogpt,
  title={BioGPT: generative pre-trained transformer for biomedical text generation and mining},
  author={Luo, Renqian and Sun, Liai and Xia, Yingce and Qin, Tao and Zhang, Sheng and Poon, Hoifung and Liu, Tie-Yan},
  journal={Briefings in bioinformatics},
  volume={23},
  number={6},
  pages={bbac409},
  year={2022},
  publisher={Oxford University Press}
}

@article{yuan2022biobart,
  title={BioBART: Pretraining and evaluation of a biomedical generative language model},
  author={Yuan, Hongyi and Yuan, Zheng and Gan, Ruyi and Zhang, Jiaxing and Xie, Yutao and Yu, Sheng},
  journal={arXiv preprint arXiv:2204.03905},
  year={2022}
}

@article{phan2021scifive,
  title={Scifive: a text-to-text transformer model for biomedical literature},
  author={Phan, Long N and Anibal, James T and Tran, Hieu and Chanana, Shaurya and Bahadroglu, Erol and Peltekian, Alec and Altan-Bonnet, Gr{\'e}goire},
  journal={arXiv preprint arXiv:2106.03598},
  year={2021}
}


@article{alsentzer2019publicly,
  title={Publicly available clinical BERT embeddings},
  author={Alsentzer, Emily and Murphy, John R and Boag, Willie and Weng, Wei-Hung and Jin, Di and Naumann, Tristan and McDermott, Matthew},
  journal={arXiv preprint arXiv:1904.03323},
  year={2019}
}


@article{rasmy2021med,
  title={Med-BERT: pretrained contextualized embeddings on large-scale structured electronic health records for disease prediction},
  author={Rasmy, Laila and Xiang, Yang and Xie, Ziqian and Tao, Cui and Zhi, Degui},
  journal={NPJ digital medicine},
  volume={4},
  number={1},
  pages={86},
  year={2021},
  publisher={Nature Publishing Group UK London}
}

@article{yang2022large,
  title={A large language model for electronic health records},
  author={Yang, Xi and Chen, Aokun and PourNejatian, Nima and Shin, Hoo Chang and Smith, Kaleb E and Parisien, Christopher and Compas, Colin and Martin, Cheryl and Costa, Anthony B and Flores, Mona G and others},
  journal={NPJ digital medicine},
  volume={5},
  number={1},
  pages={194},
  year={2022},
  publisher={Nature Publishing Group UK London}
}

@inproceedings{ouyang2022training,
  title={Training language models to follow instructions with human feedback},
  author={Ouyang, Long and Wu, Jeff and Jiang, Xu and Almeida, Diogo and Wainwright, Carroll L and Mishkin, Pamela and Zhang, Chong and Agarwal, Sandhini and Slama, Katarina and Ray, Alex and Schulman, John and Hilton, Jacob and Kelton, Fraser and Miller, Luke and Simens, Maddie and Askell, Amanda and Welinder, Peter and Christiano, Paul and Leike, Jan and Lowe, Ryan},
  booktitle={Advances in Neural Information Processing Systems},
  volume={35},
  pages={27730--27744},
  year={2022},
  url={https://proceedings.neurips.cc/paper_files/paper/2022/file/b1efde53be364a73914f58805a001731-Paper-Conference.pdf}
}


@inproceedings{christiano2017deep,
  title={Deep reinforcement learning from human preferences},
  author={Christiano, Paul F and Leike, Jan and Brown, Tom B and Martic, Miljan and Legg, Shane and Amodei, Dario},
  booktitle={Advances in Neural Information Processing Systems},
  volume={30},
  pages={4299--4307},
  year={2017},
  url={https://proceedings.neurips.cc/paper_files/paper/2017/file/d5e2c0adad503c91f91df240d0cd4e49-Paper.pdf}
}

@inproceedings{stiennon2020learning,
  title={Learning to summarize from human feedback},
  author={Stiennon, Nisan and Ouyang, Long and Wu, Jeff and Ziegler, Daniel M and Lowe, Ryan and Voss, Chelsea and Radford, Alec and Amodei, Dario and Christiano, Paul},
  booktitle={Advances in Neural Information Processing Systems},
  volume={33},
  pages={3008--3021},
  year={2020},
  url={https://proceedings.neurips.cc/paper_files/paper/2020/file/1f89885d556929e98d3ef9b86448f951-Paper.pdf}
}

@inproceedings{rafailov2023direct,
  title={Direct Preference Optimization: Your Language Model is Secretly a Reward Model},
  author={Rafailov, Rafael and Sharma, Archit and Mitchell, Eric and Ermon, Stefano and Manning, Christopher D and Finn, Chelsea},
  booktitle={Advances in Neural Information Processing Systems},
  volume={36},
  year={2023},
  url={https://openreview.net/pdf?id=HPuSIXJaa9}
}

@article{yin2024relative,
  title={Relative Preference Optimization: Enhancing LLM Alignment through Contrasting Responses},
  author={Yin, Yueqin and others},
  journal={arXiv preprint arXiv:2402.10958},
  year={2024},
  url={https://arxiv.org/abs/2402.10958}
}

@article{kiruluta2025self,
  title={A Self-Supervised Reinforcement Learning Approach for Fine-Tuning Large Language Models Using Cross-Attention Signals},
  author={Kiruluta, Andrew and Lemos, Andreas and Burity, Priscilla},
  journal={arXiv preprint arXiv:2502.10482},
  year={2025},
  url={https://arxiv.org/abs/2502.10482}
}


@article{yin2025genome,
  title={Genome-Bench: A Scientific Reasoning Benchmark from Real-World CRISPR Discussions},
  author={Anonymous Authors},
  journal={},
  year={2025},
}


@article{xiong2025minimalist,
  title={A Minimalist Approach to LLM Reasoning: From Rejection Sampling to Reinforce},
  author={Xiong, Wei and Yao, Jiarui and Xu, Yuhui and Pang, Bo and Wang, Lei and Sahoo, Doyen and Li, Junnan and Jiang, Nan and Zhang, Tong and Xiong, Caiming and Dong, Hanze},
  journal={arXiv preprint arXiv:2504.11343},
  year={2025},
  url={https://arxiv.org/abs/2504.11343}
}

@article{lambert2024t,
  title={T$\backslash$" ulu 3: Pushing frontiers in open language model post-training},
  author={Lambert, Nathan and Morrison, Jacob and Pyatkin, Valentina and Huang, Shengyi and Ivison, Hamish and Brahman, Faeze and Miranda, Lester James V and Liu, Alisa and Dziri, Nouha and Lyu, Shane and others},
  journal={arXiv preprint arXiv:2411.15124},
  year={2024}
}

@article{liu2025understanding,
  title={Understanding r1-zero-like training: A critical perspective},
  author={Liu, Zichen and Chen, Changyu and Li, Wenjun and Qi, Penghui and Pang, Tianyu and Du, Chao and Lee, Wee Sun and Lin, Min},
  journal={arXiv preprint arXiv:2503.20783},
  year={2025}
}


@inproceedings{rein2024gpqa,
  title={Gpqa: A graduate-level google-proof q\&a benchmark},
  author={Rein, David and Hou, Betty Li and Stickland, Asa Cooper and Petty, Jackson and Pang, Richard Yuanzhe and Dirani, Julien and Michael, Julian and Bowman, Samuel R},
  booktitle={First Conference on Language Modeling},
  year={2024}
}

@article{phan2025humanity,
  title={Humanity's Last Exam},
  author={Phan, Long and Gatti, Alice and Han, Ziwen and Li, Nathaniel and Hu, Josephina and Zhang, Hugh and Zhang, Chen Bo Calvin and Shaaban, Mohamed and Ling, John and Shi, Sean and others},
  journal={arXiv preprint arXiv:2501.14249},
  year={2025}
}

@article{laurent2024lab,
  title={Lab-bench: Measuring capabilities of language models for biology research},
  author={Laurent, Jon M and Janizek, Joseph D and Ruzo, Michael and Hinks, Michaela M and Hammerling, Michael J and Narayanan, Siddharth and Ponnapati, Manvitha and White, Andrew D and Rodriques, Samuel G},
  journal={arXiv preprint arXiv:2407.10362},
  year={2024}
}

@article{chu20245,
  title={A 5′ UTR language model for decoding untranslated regions of mRNA and function predictions},
  author={Chu, Yanyi and Yu, Dan and Li, Yupeng and Huang, Kaixuan and Shen, Yue and Cong, Le and Zhang, Jason and Wang, Mengdi},
  journal={Nature Machine Intelligence},
  volume={6},
  number={4},
  pages={449--460},
  year={2024},
  publisher={Nature Publishing Group UK London}
}

@article{guo2024embodied,
  title={Embodied llm agents learn to cooperate in organized teams},
  author={Guo, Xudong and Huang, Kaixuan and Liu, Jiale and Fan, Wenhui and V{\'e}lez, Natalia and Wu, Qingyun and Wang, Huazheng and Griffiths, Thomas L and Wang, Mengdi},
  journal={arXiv preprint arXiv:2403.12482},
  year={2024}
}

@article{wang2025call,
  title={A call for built-in biosecurity safeguards for generative AI tools},
  author={Wang, Mengdi and Zhang, Zaixi and Bedi, Amrit Singh and Velasquez, Alvaro and Guerra, Stephanie and Lin-Gibson, Sheng and Cong, Le and Qu, Yuanhao and Chakraborty, Souradip and Blewett, Megan and others},
  journal={Nature Biotechnology},
  pages={1--3},
  year={2025},
  publisher={Nature Publishing Group}
}


@article{guo2025temporal,
  title={Temporal Consistency for LLM Reasoning Process Error Identification},
  author={Guo, Jiacheng and Wu, Yue and Qiu, Jiahao and Huang, Kaixuan and Juan, Xinzhe and Yang, Ling and Wang, Mengdi},
  journal={arXiv preprint arXiv:2503.14495},
  year={2025}
}

@article{huang2025math,
  title={MATH-Perturb: Benchmarking LLMs' Math Reasoning Abilities against Hard Perturbations},
  author={Huang, Kaixuan and Guo, Jiacheng and Li, Zihao and Ji, Xiang and Ge, Jiawei and Li, Wenzhe and Guo, Yingqing and Cai, Tianle and Yuan, Hui and Wang, Runzhe and others},
  journal={International Conference on Machine Learning},
  year={2025}
}


@article{chakraborty2024maxmin,
  title={MaxMin-RLHF: Alignment with diverse human preferences},
  author={Chakraborty, Souradip and Qiu, Jiahao and Yuan, Hui and Koppel, Alec and Huang, Furong and Manocha, Dinesh and Bedi, Amrit Singh and Wang, Mengdi},
  journal={arXiv preprint arXiv:2402.08925},
  year={2024}
}
