## Supplementary figures and images for "Genome-Bench: A Scientific Reasoning Benchmark from Real-World Expert Discussions"

### answer_distribution.png

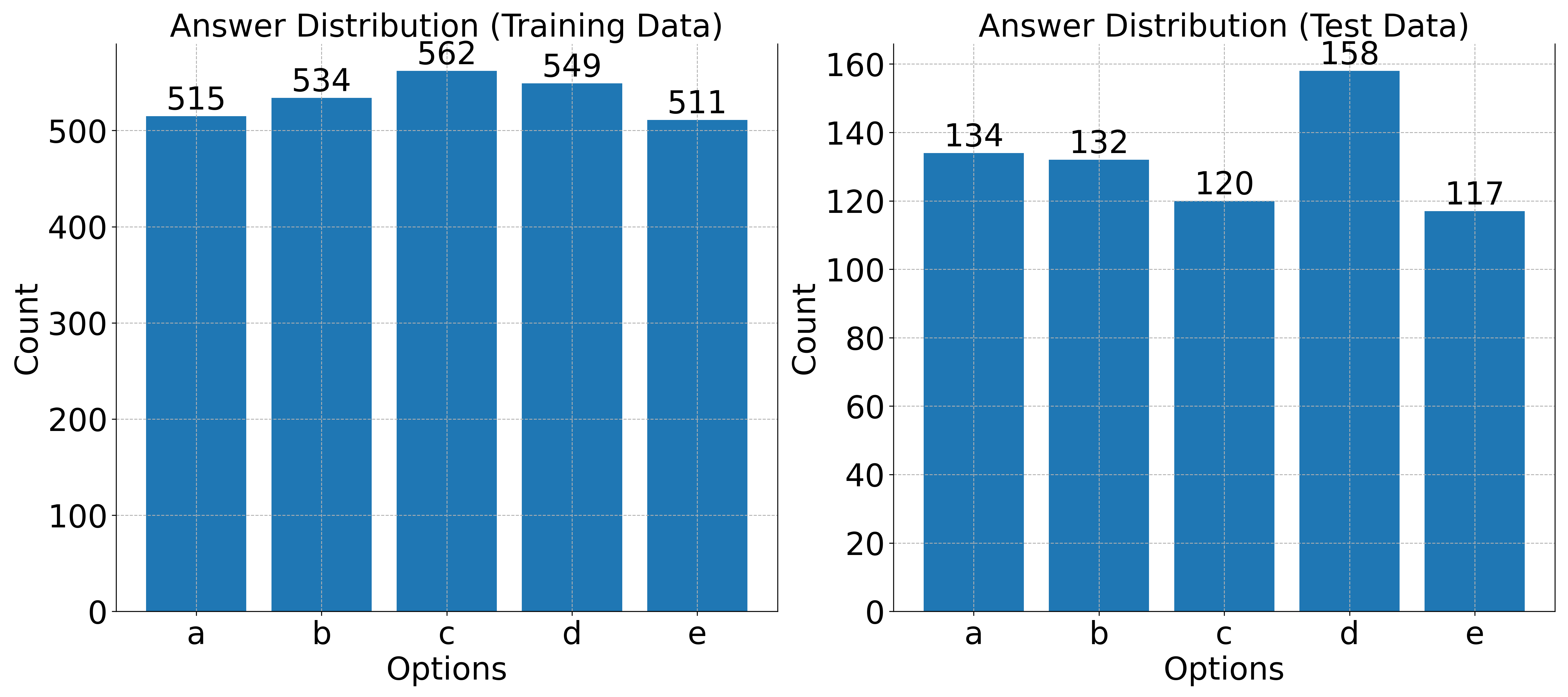

### appendix_1_2.pdf

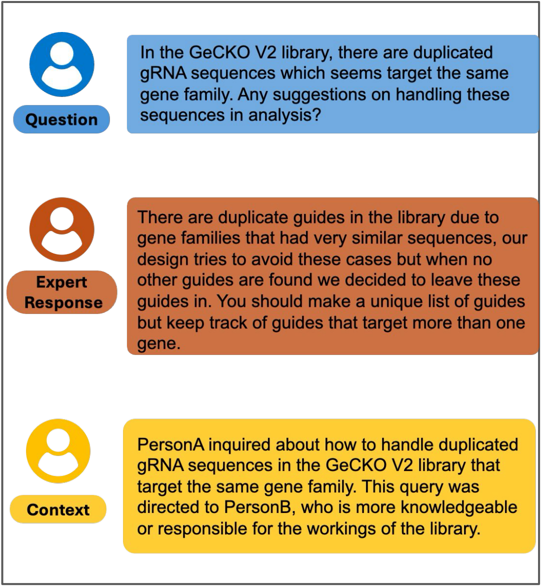

### crispr_difficulty_accuracy_barplot_api_grouped_by_model_with_gpt41.png

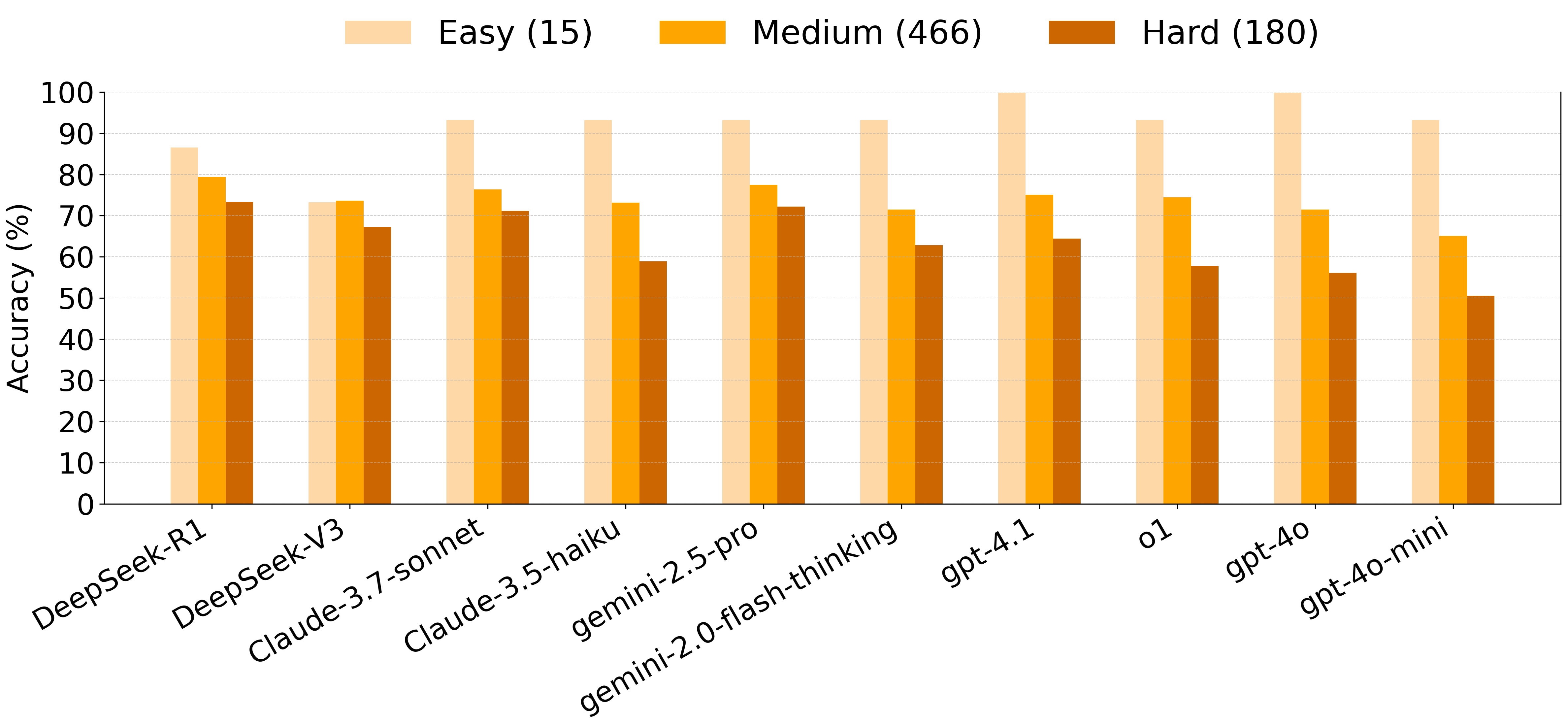

### crispr_difficulty_accuracy_barplot_local_grouped_by_model_with_gpt41.png

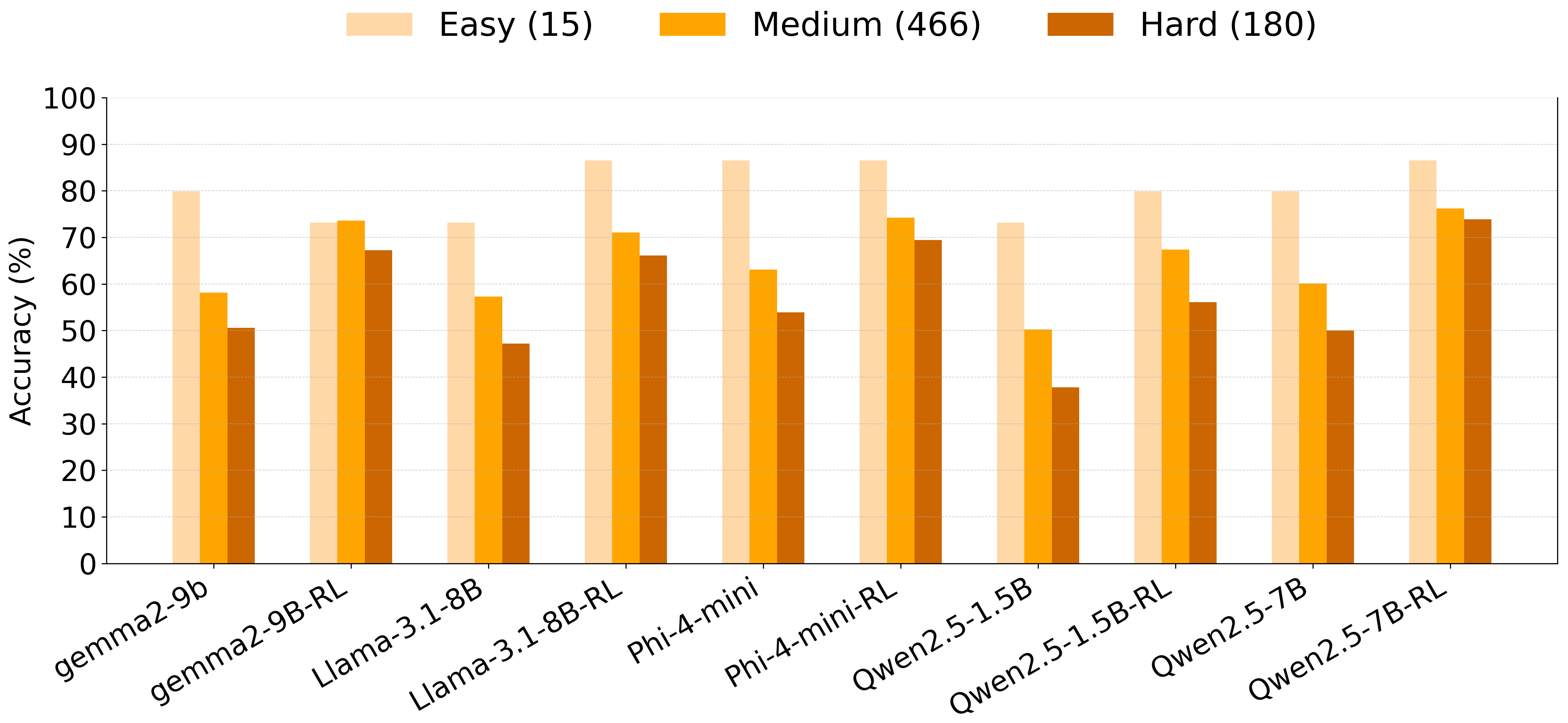

### crispr_model_grouped_accuracy_barplot_with_gpt41.png

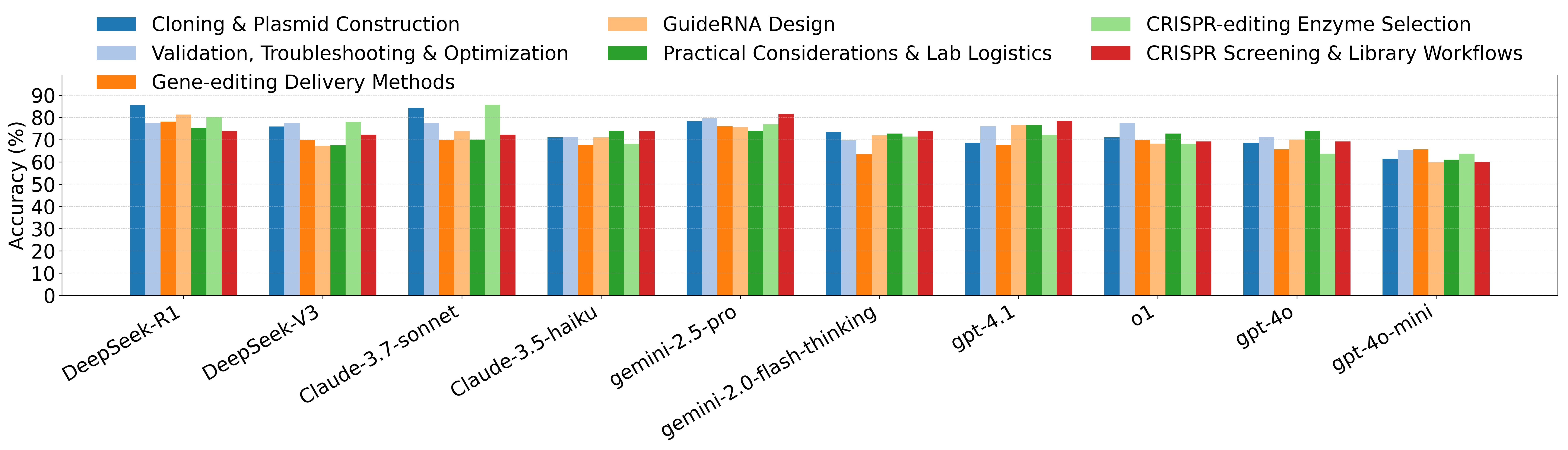

### crispr_model_grouped_accuracy_barplot_with_gpt41_local.png

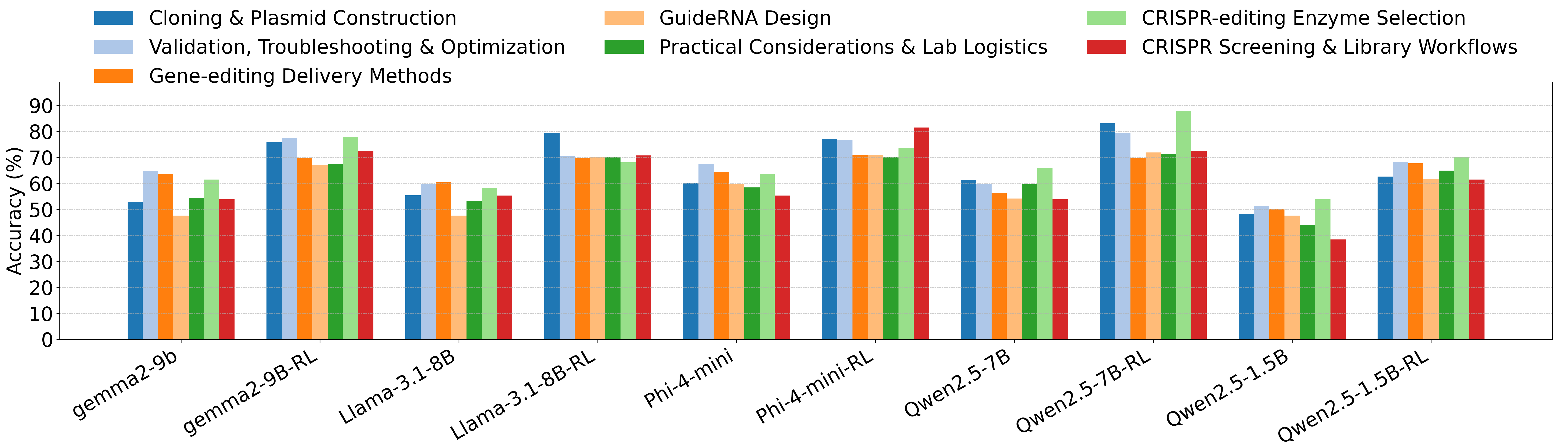

### Figure_1.pdf

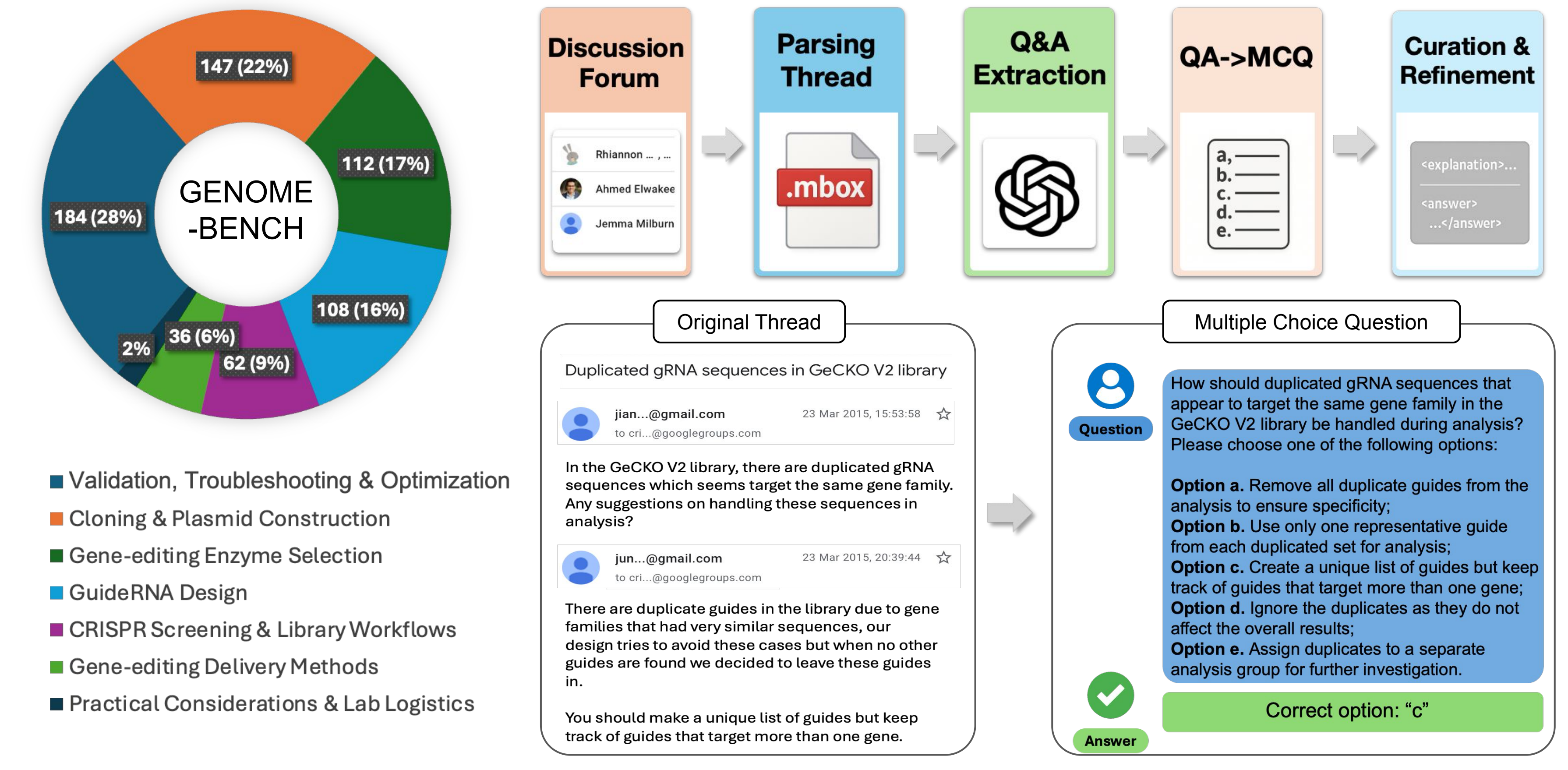

### Figure_2.pdf

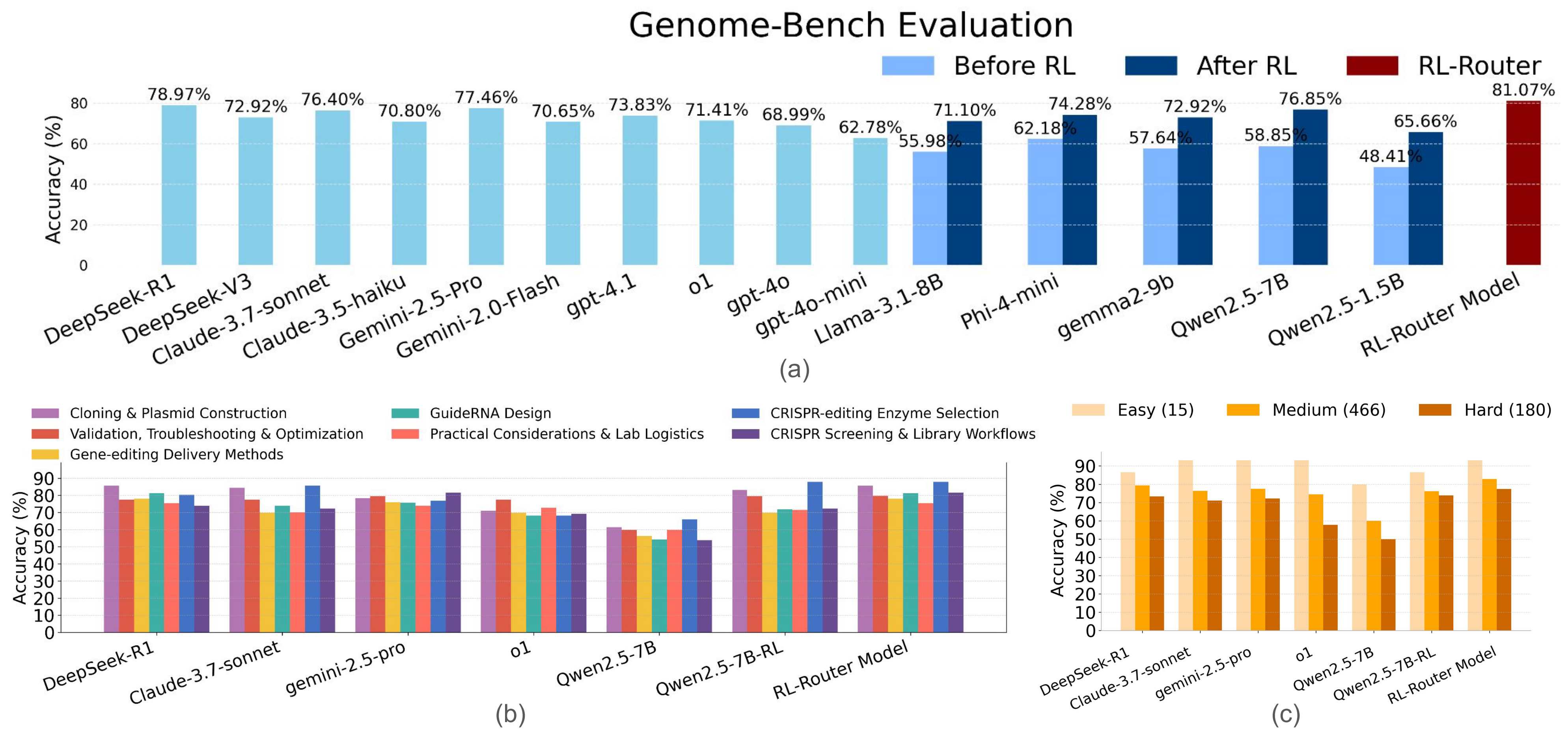

### Figure_3.pdf

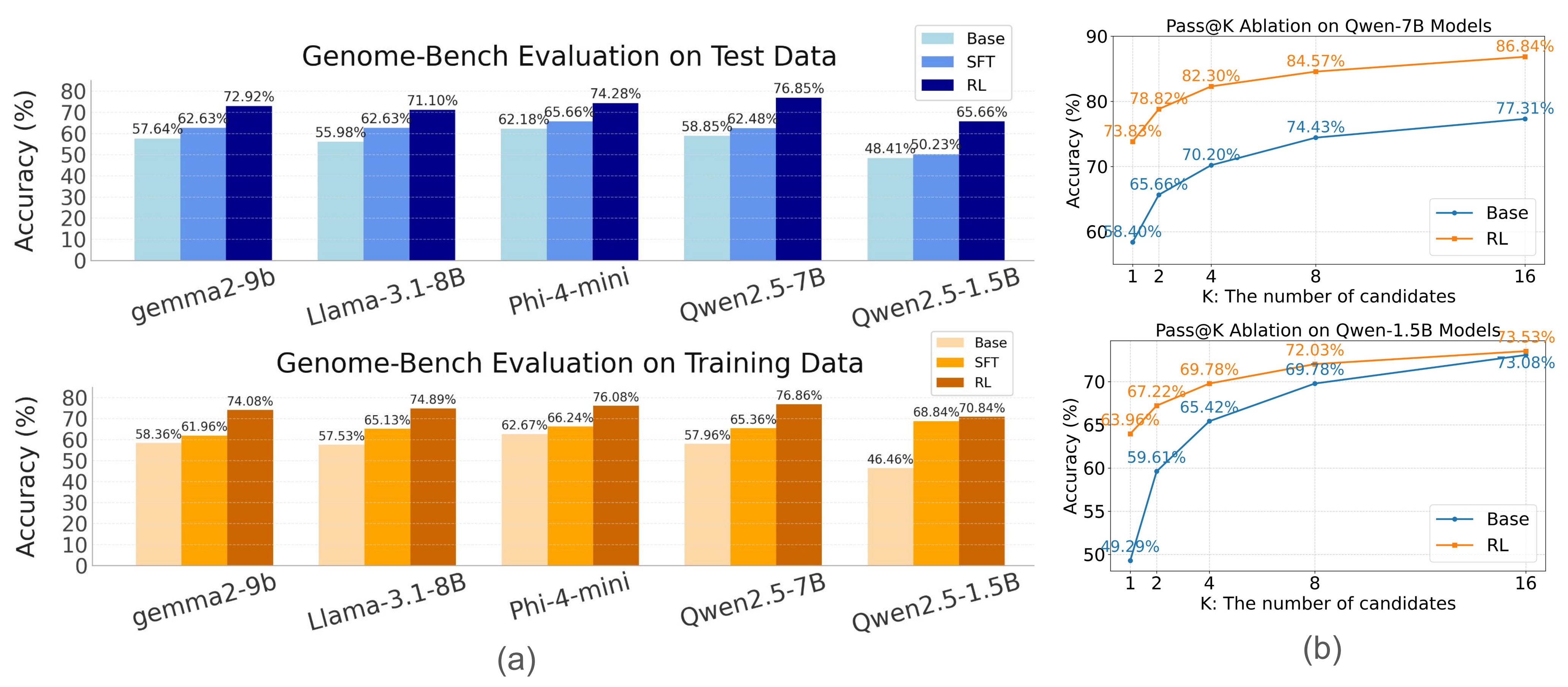

### Figure_4.pdf

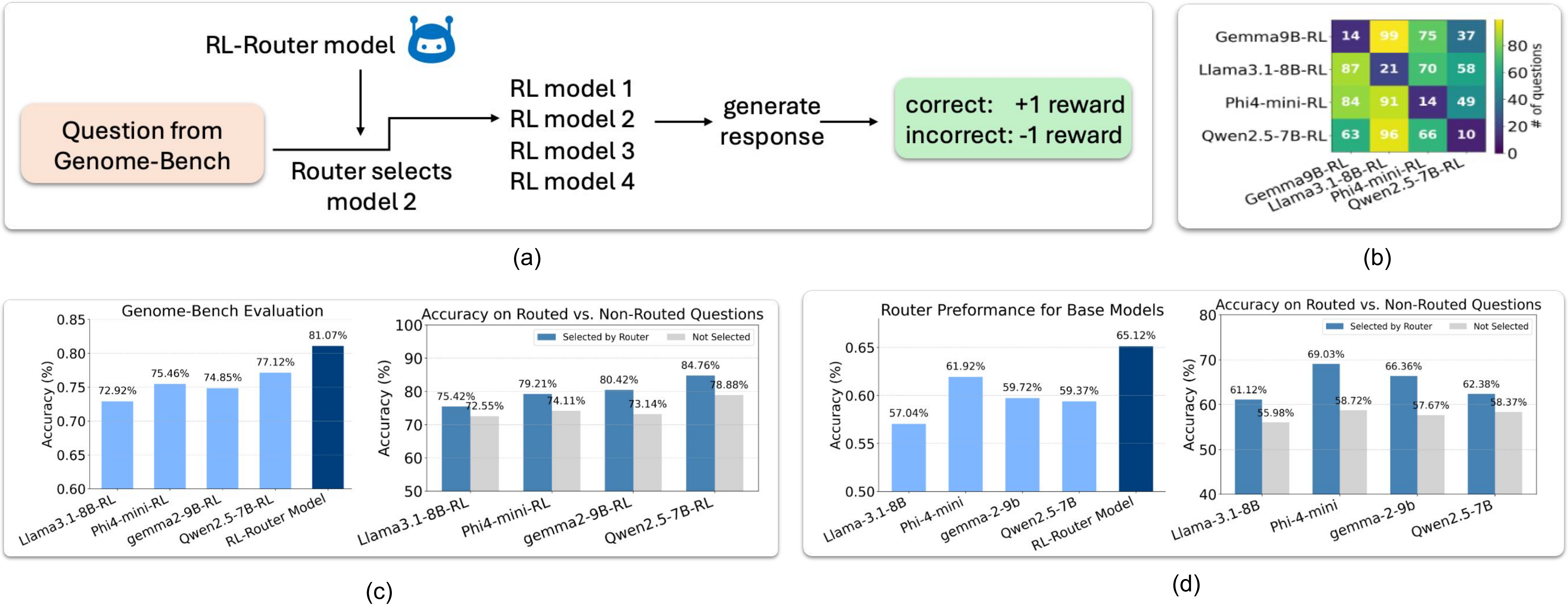

### Figure_5.pdf

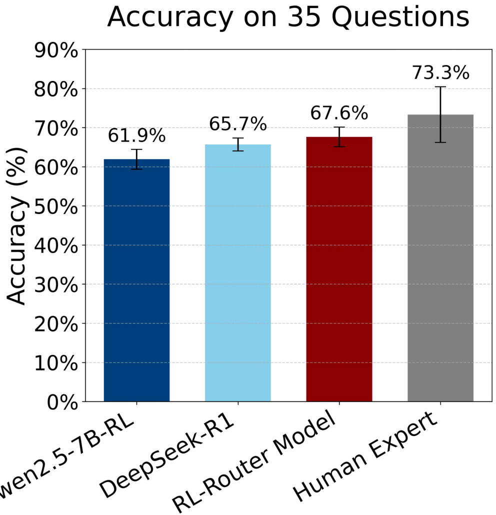

### human_expert.png

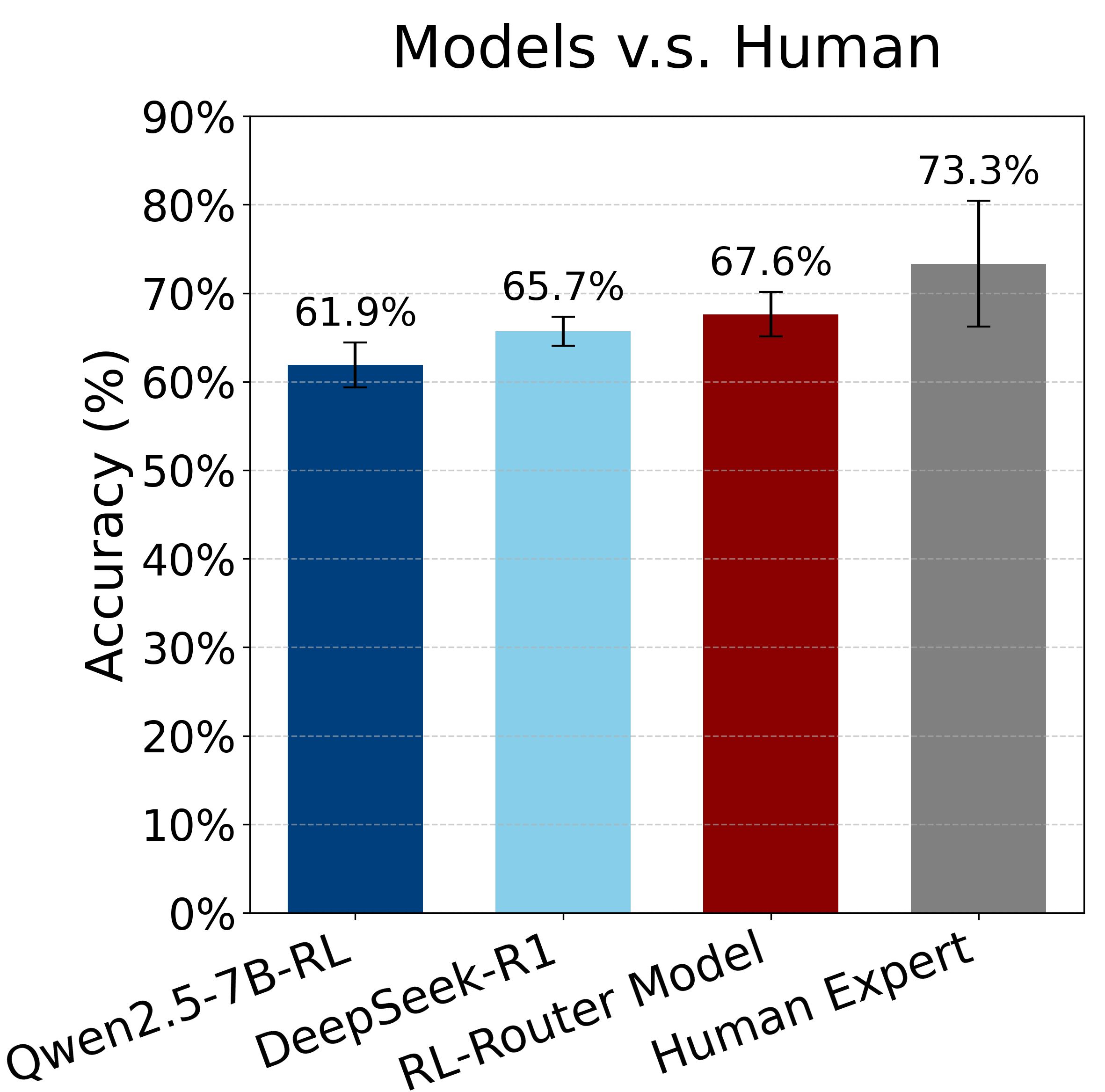

### model_accuracy_comparison_1.5B.png

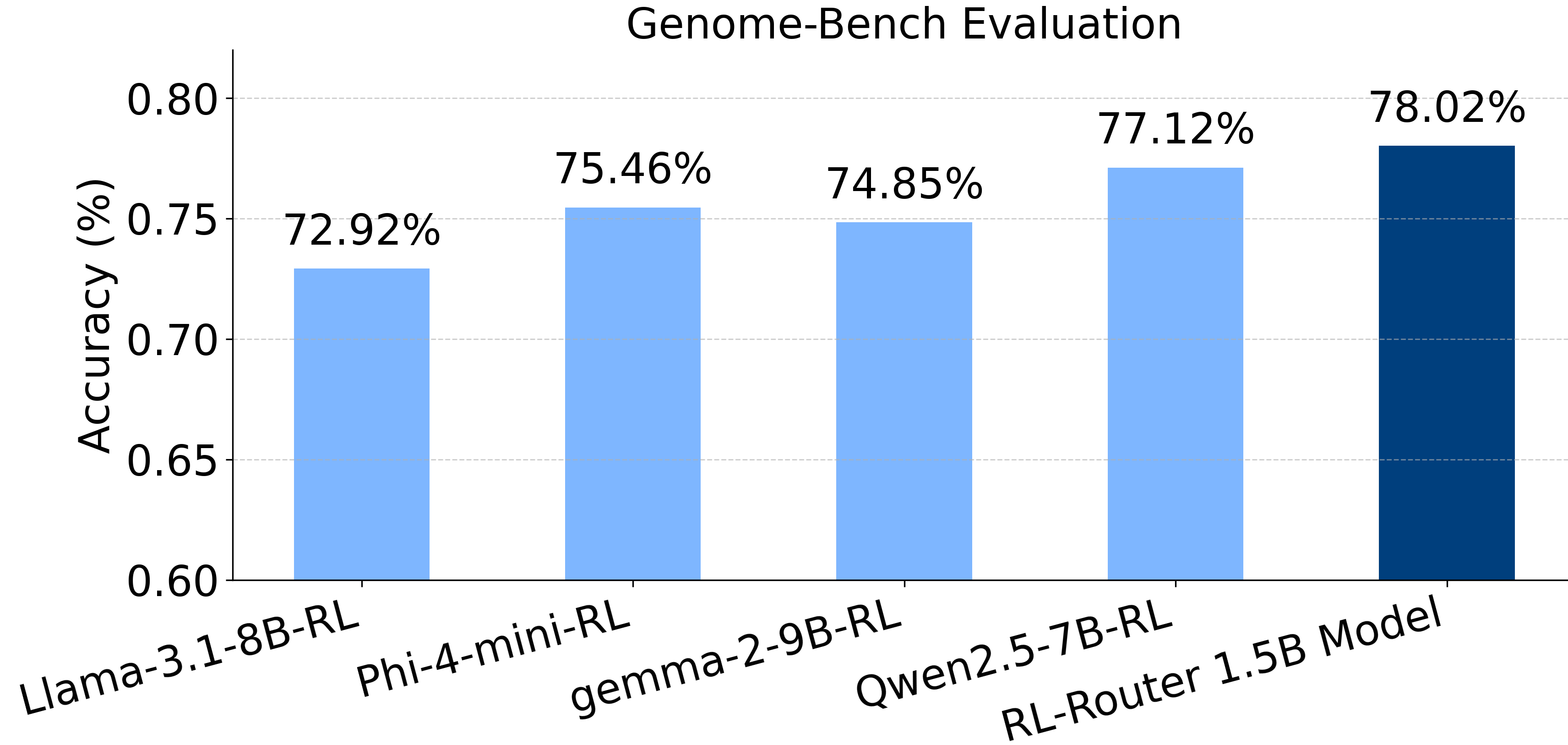

### page_1.png

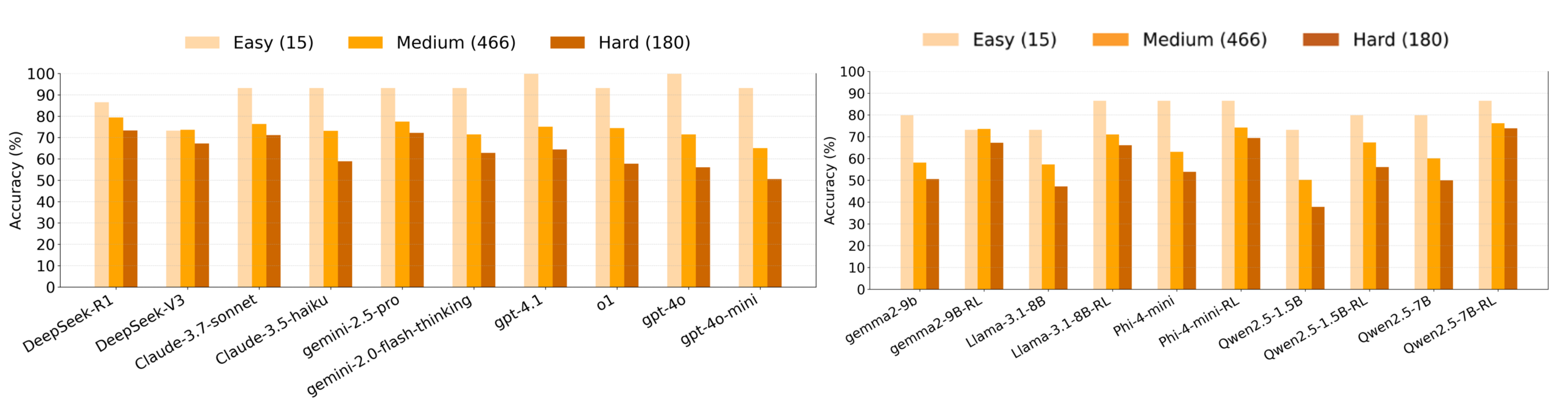

### pass_at_k_crispr_Gemma-2-9B.png

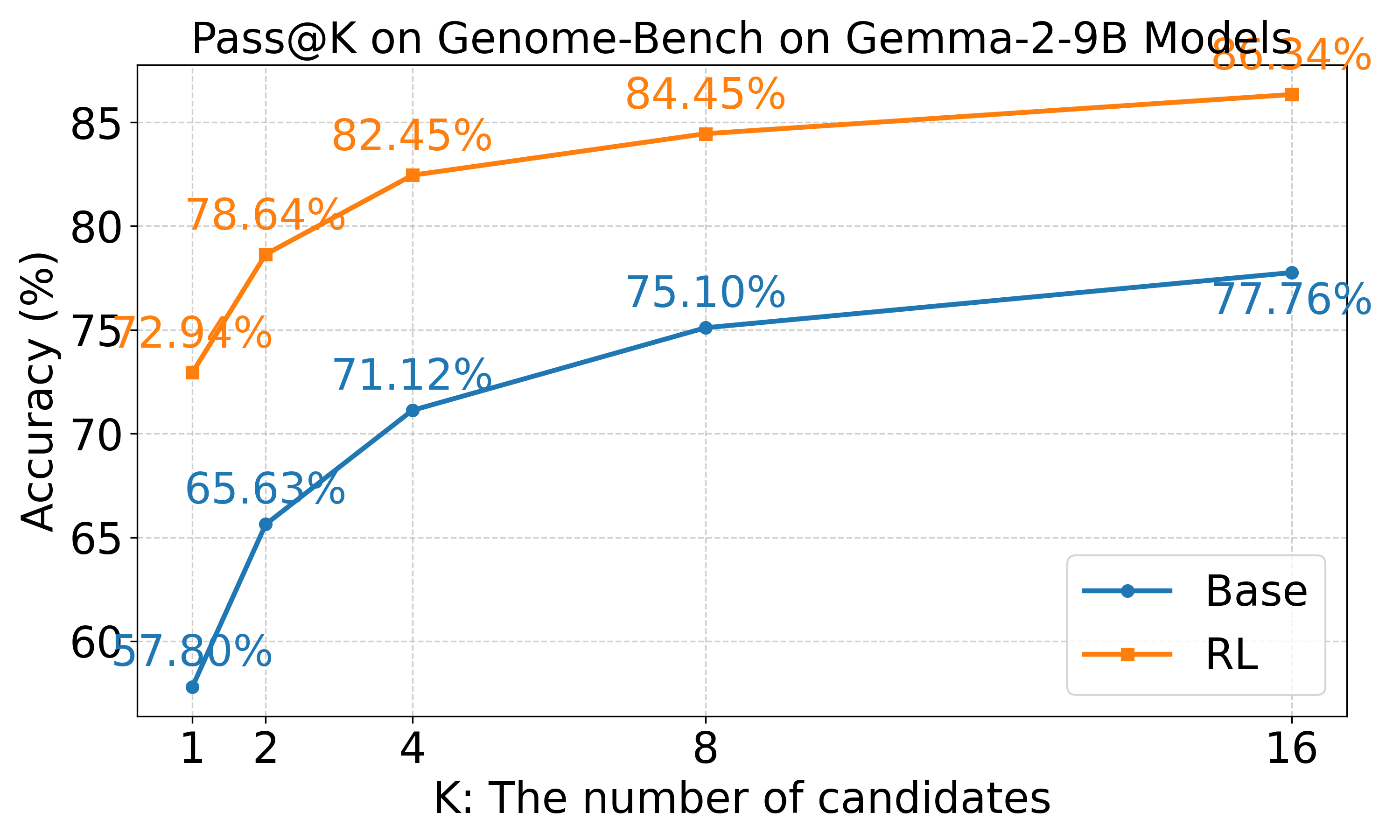

### pass_at_k_crispr_Llama-3.1-8B.png

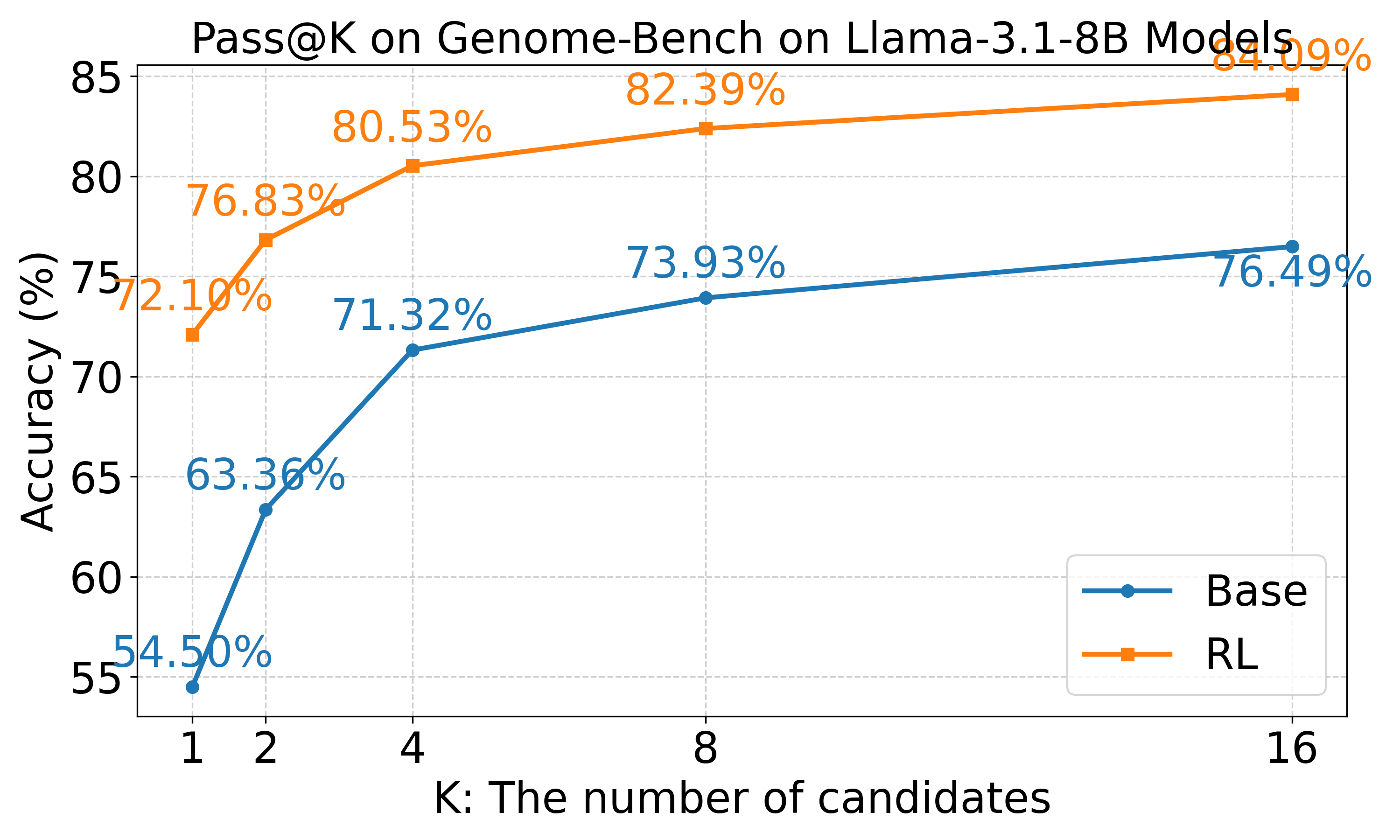

### pass_at_k_crispr_Phi-4-mini.png

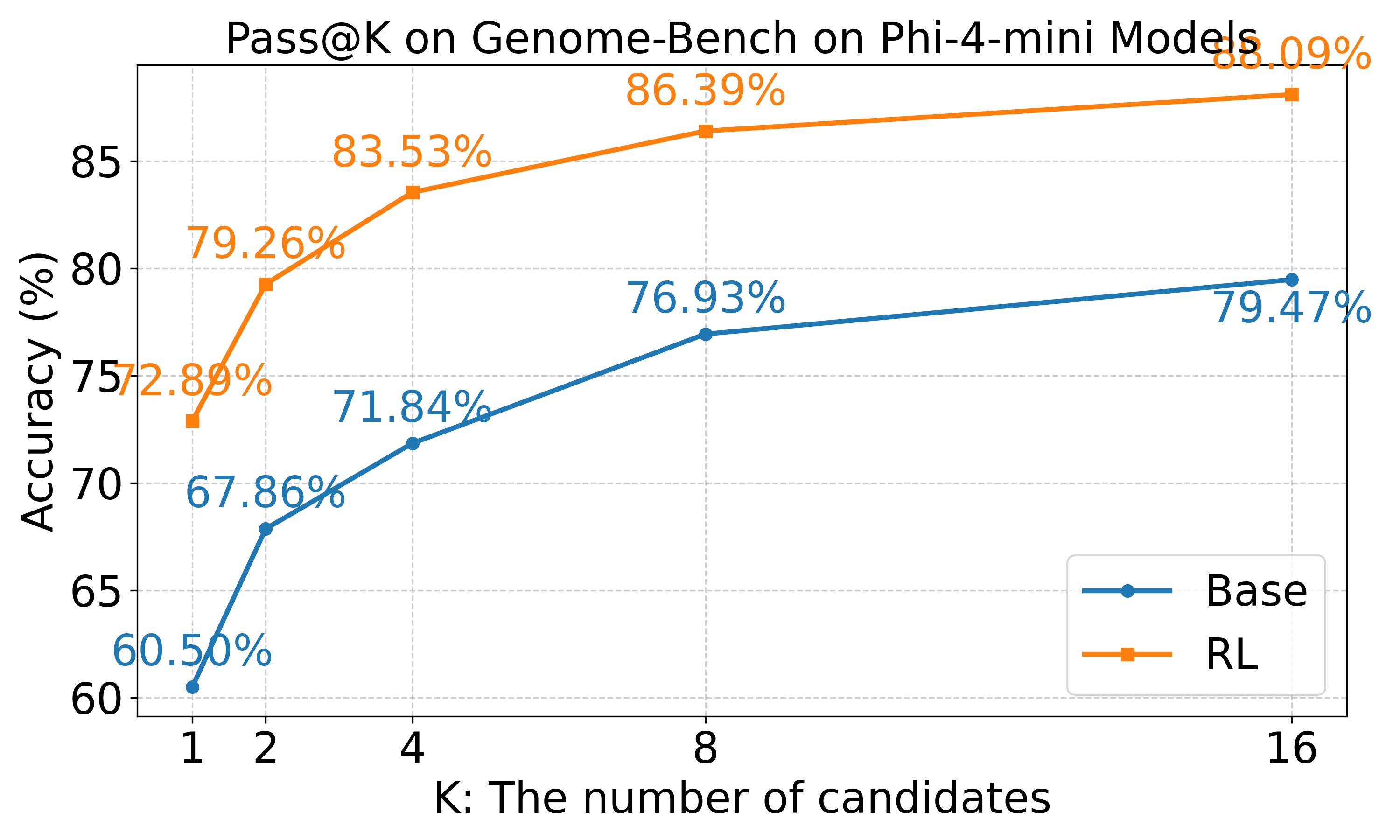

### pass_at_k_crispr_Qwen2.5-1.5B.png

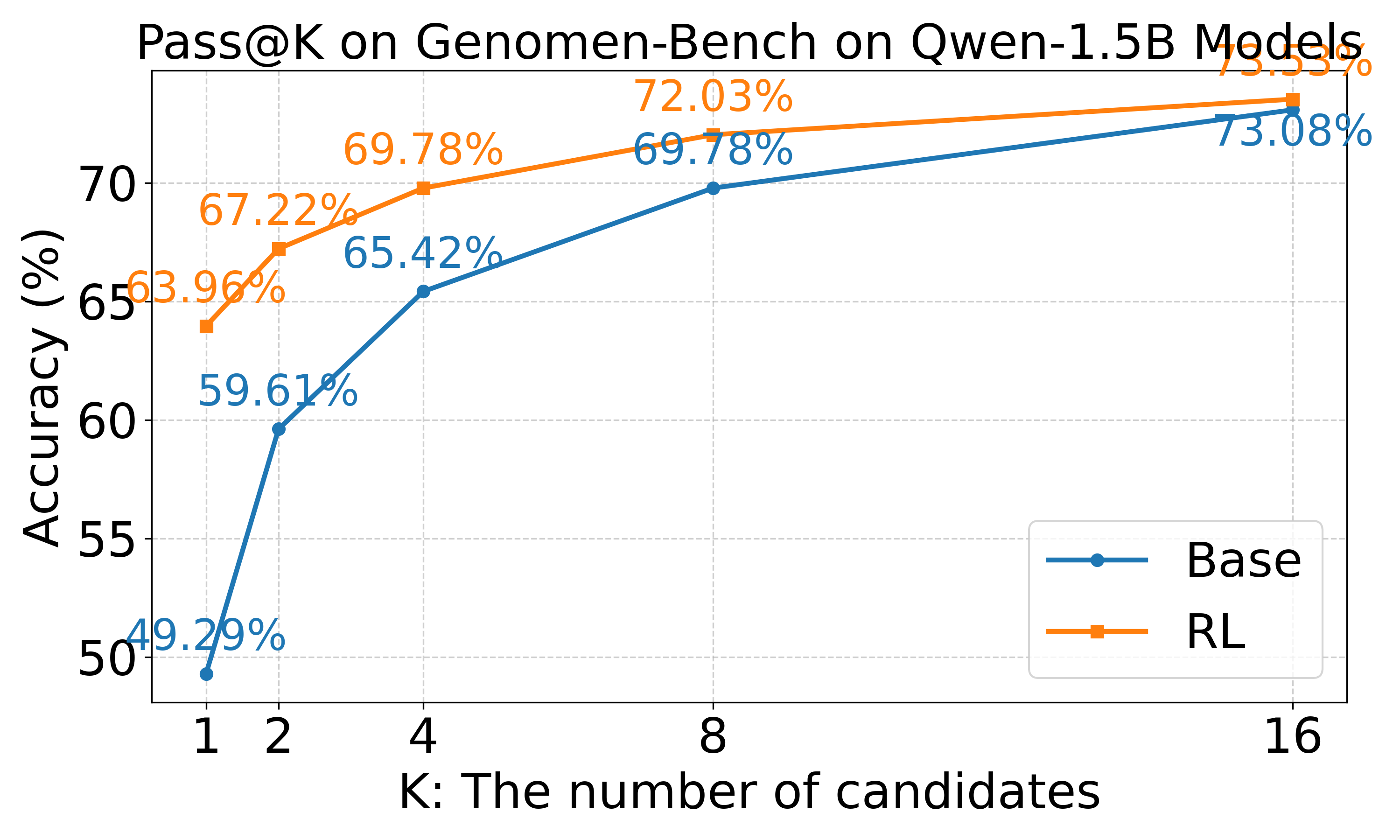

### pass_at_k_crispr_Qwen2.5-7B.png

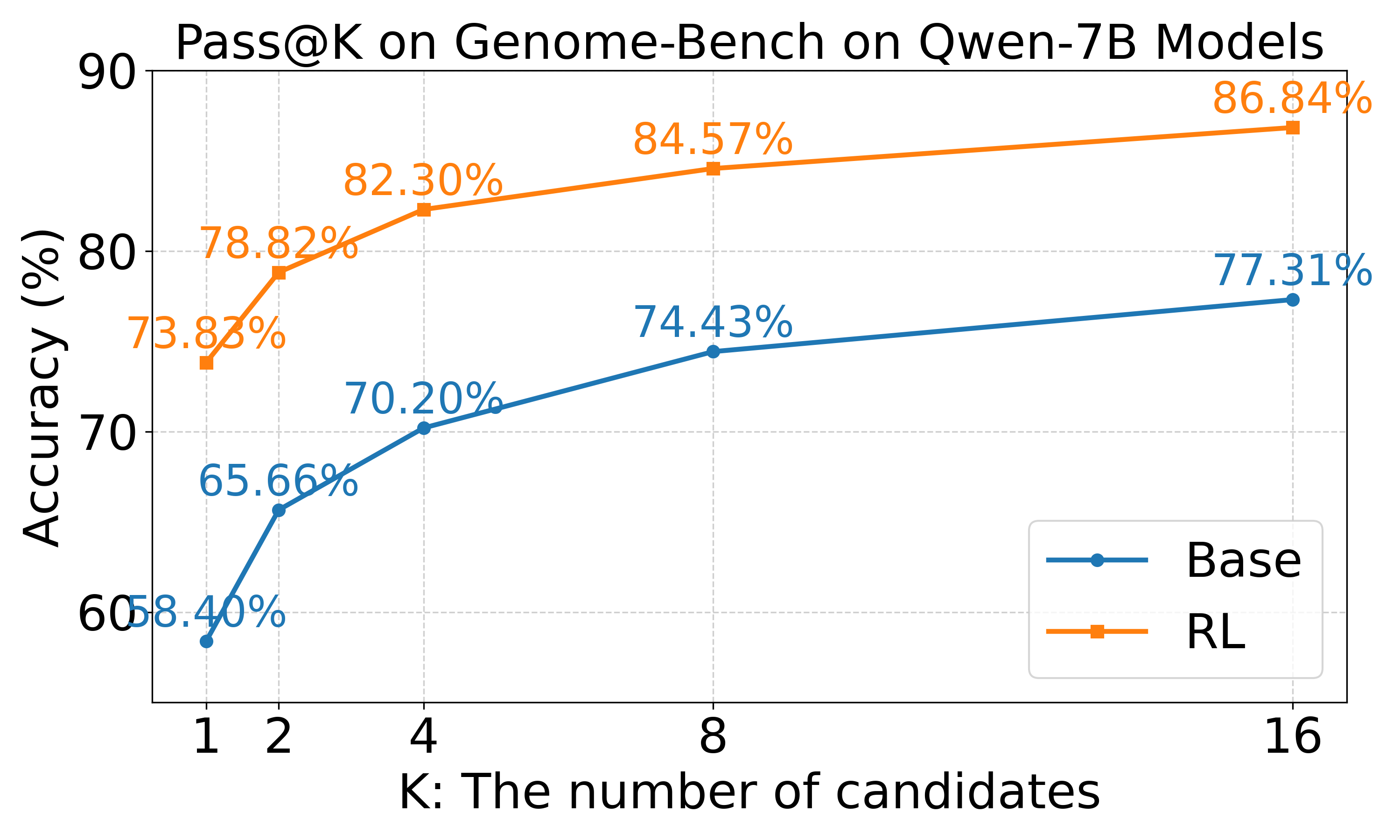

### router_distribution_1_7B.png

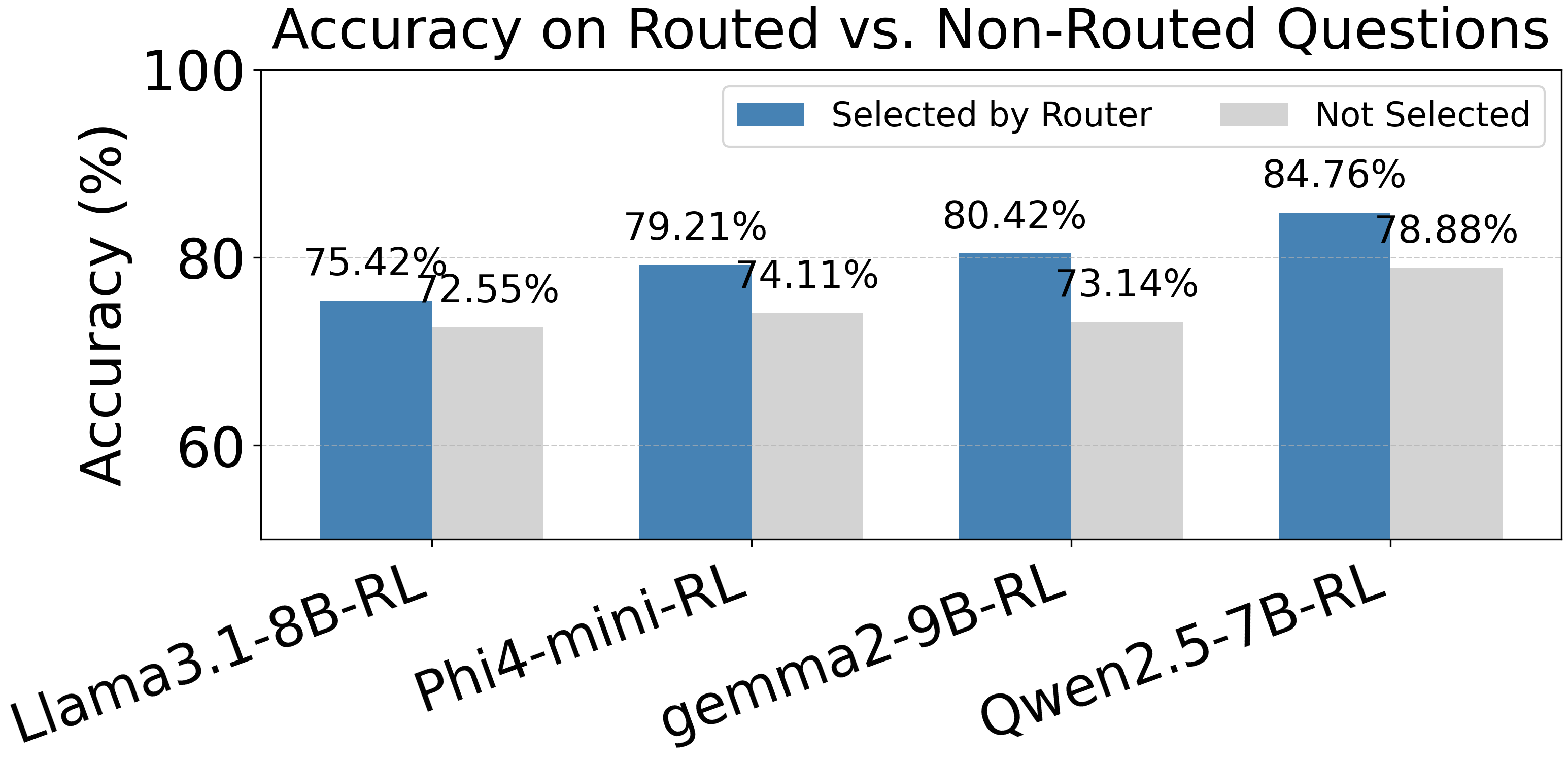

### router_distribution_2_7B.png

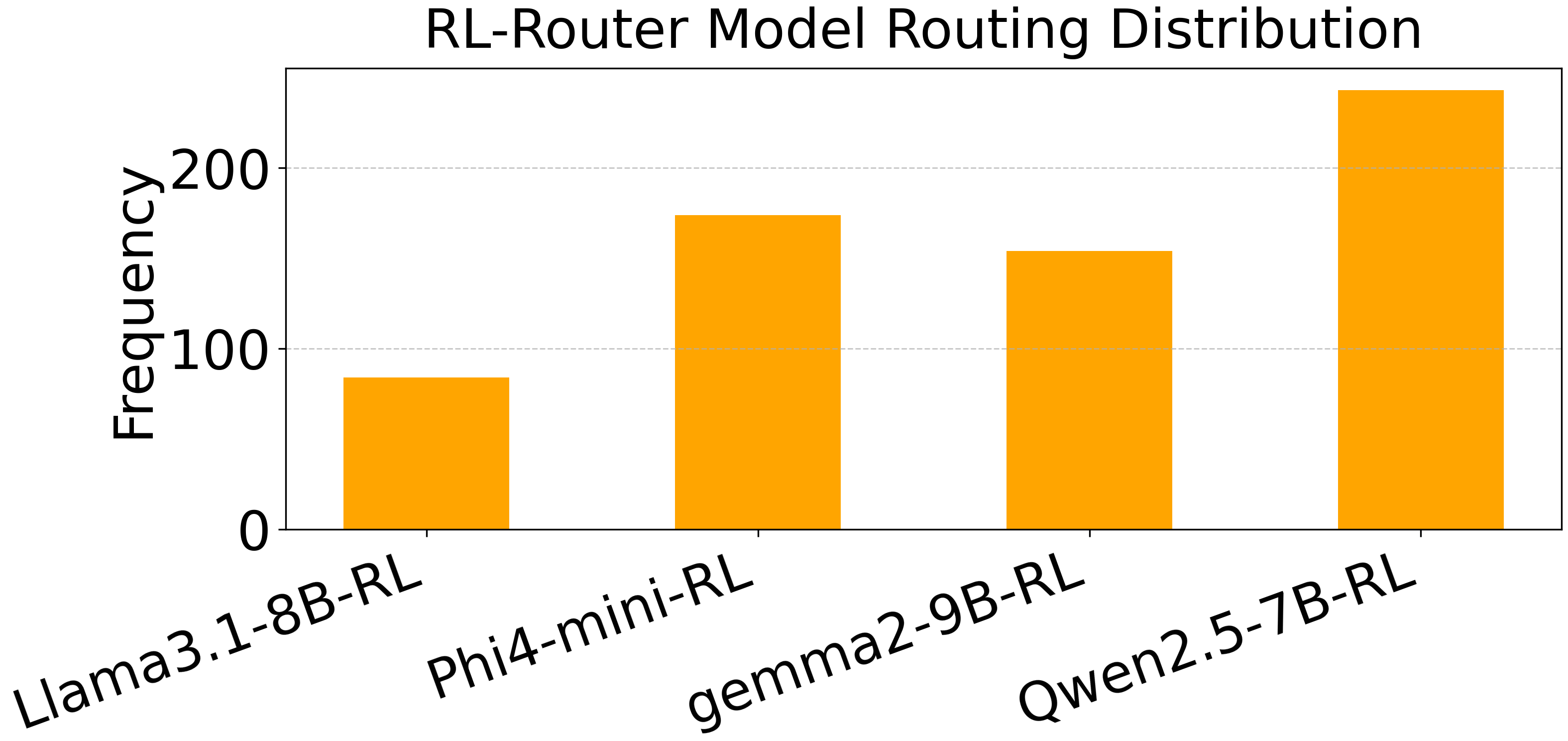

### router_distribution_7B.png

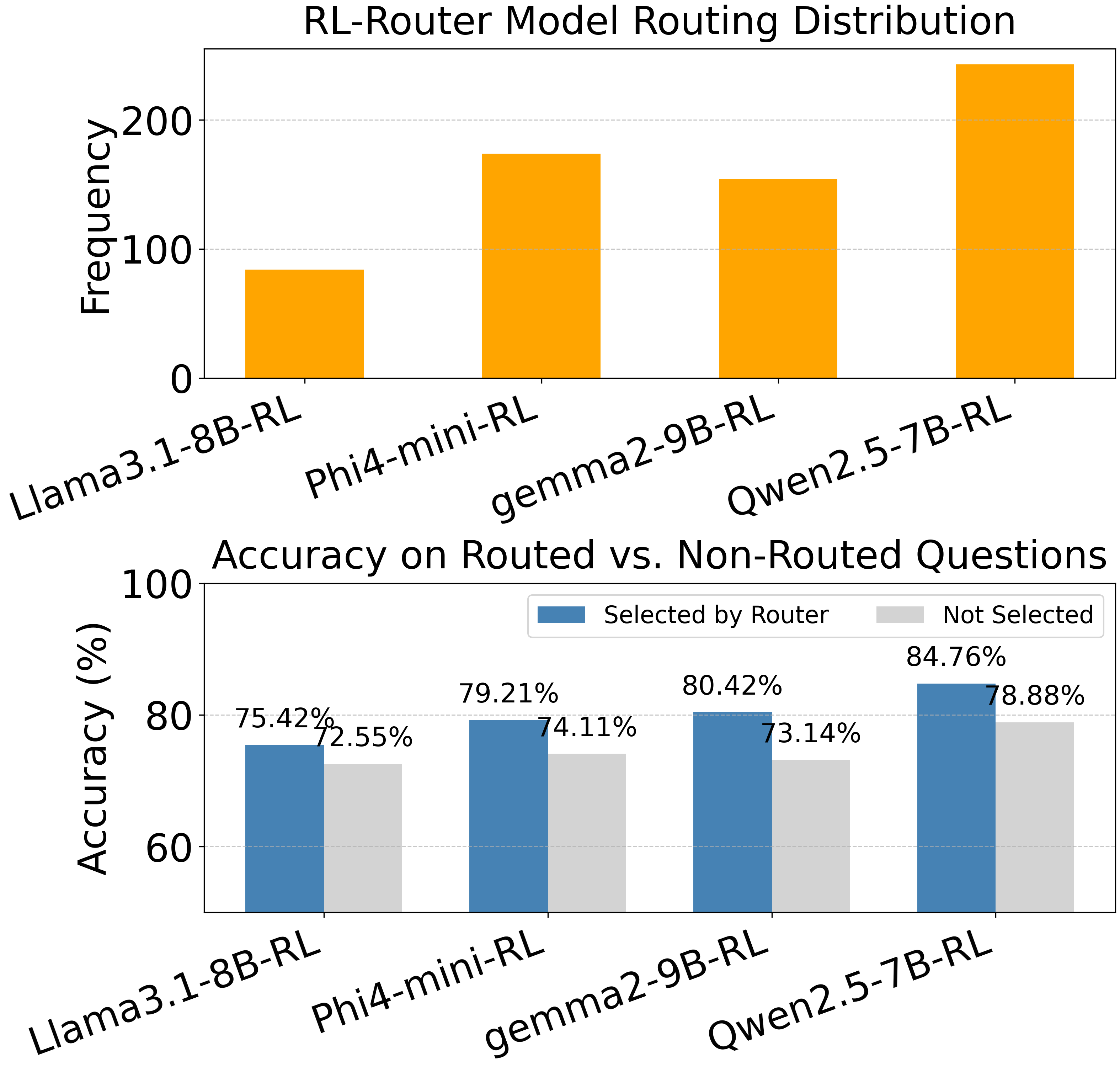

### router_distribution_and_comparison_accuracy_1.5B.png

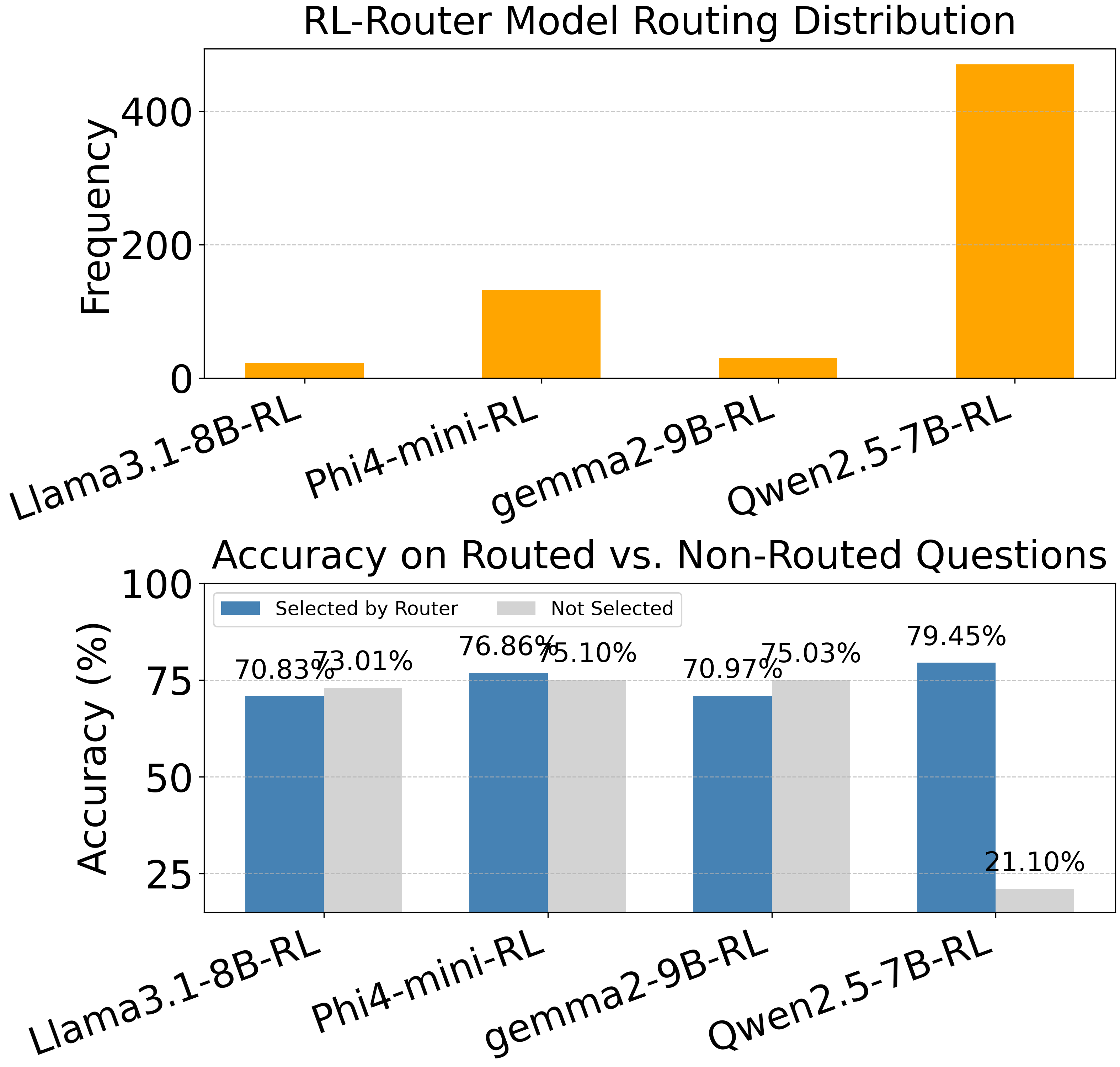

### router_distribution_no_1_7B.png

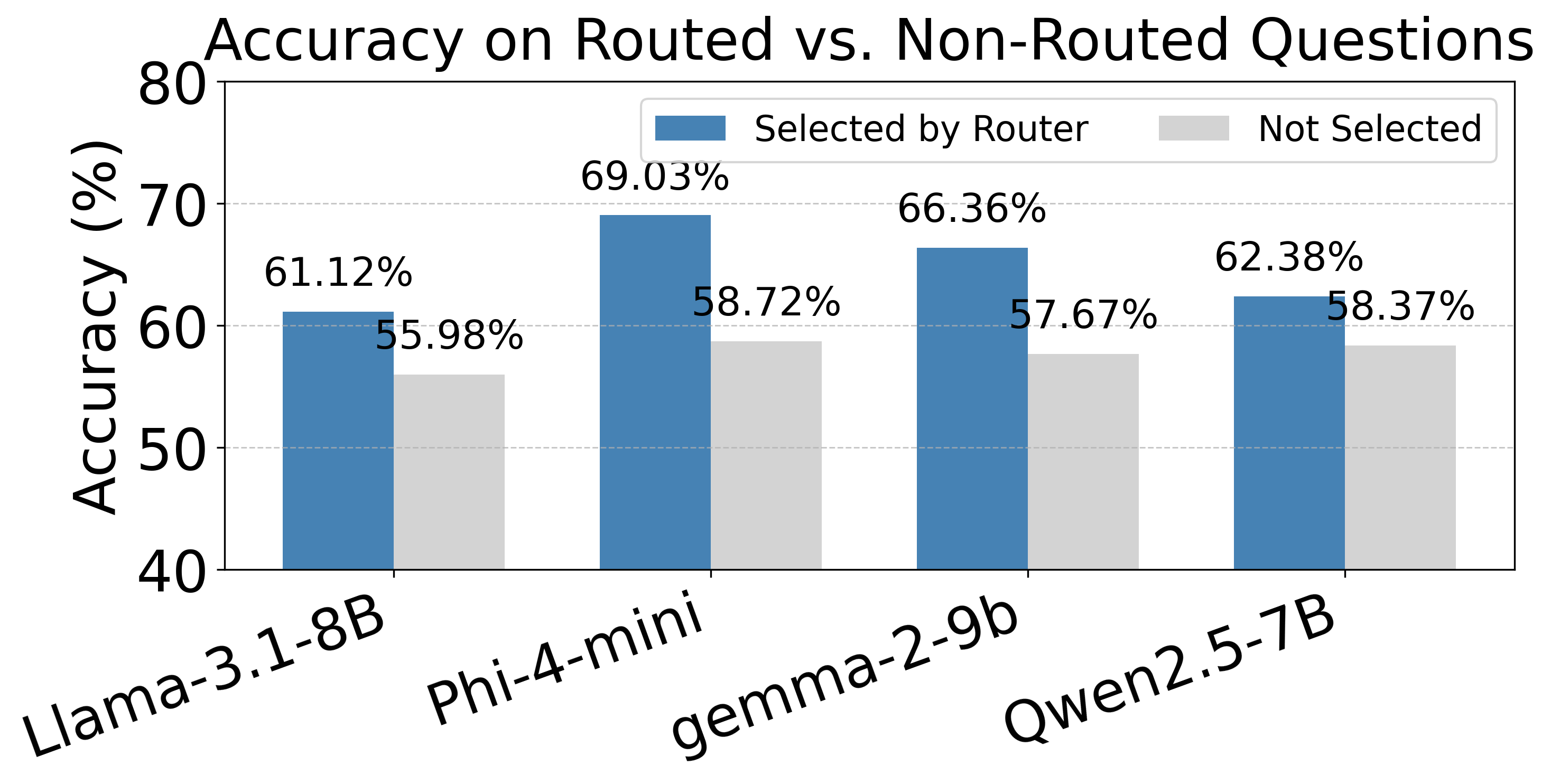

### router_distribution_no_2_7B.png

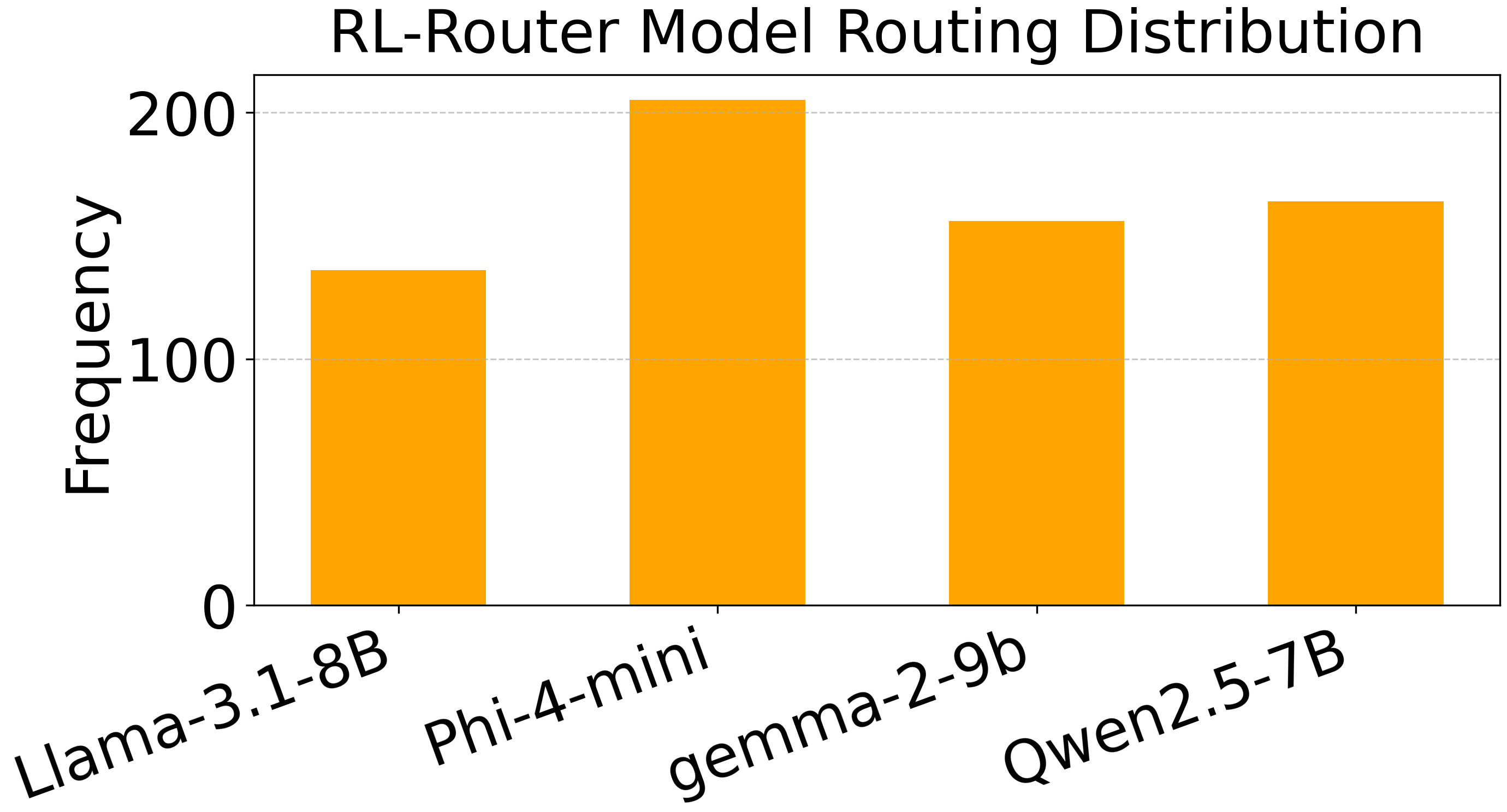

### router_distribution_no_7B.png

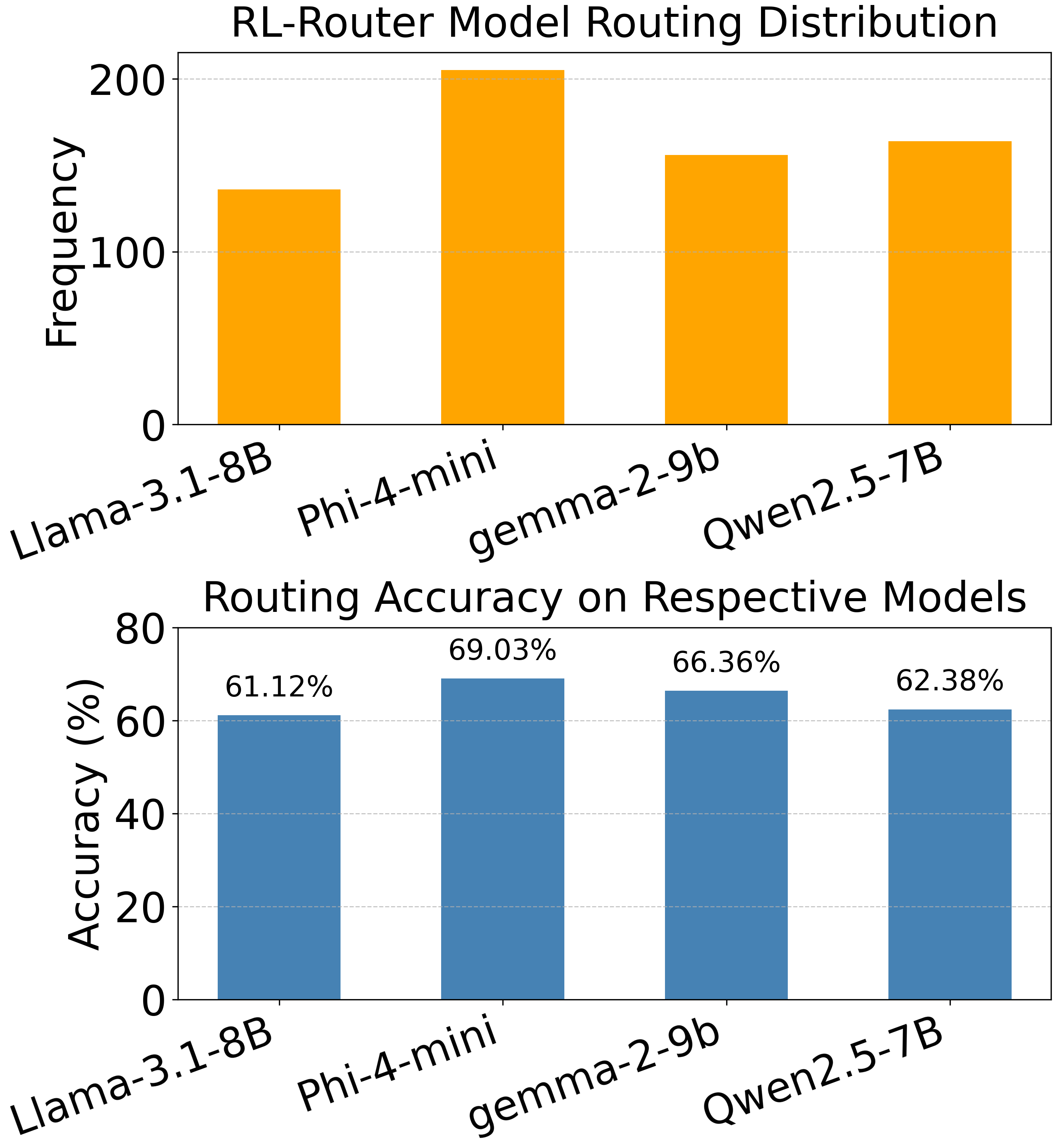

### router_performance_7B.png

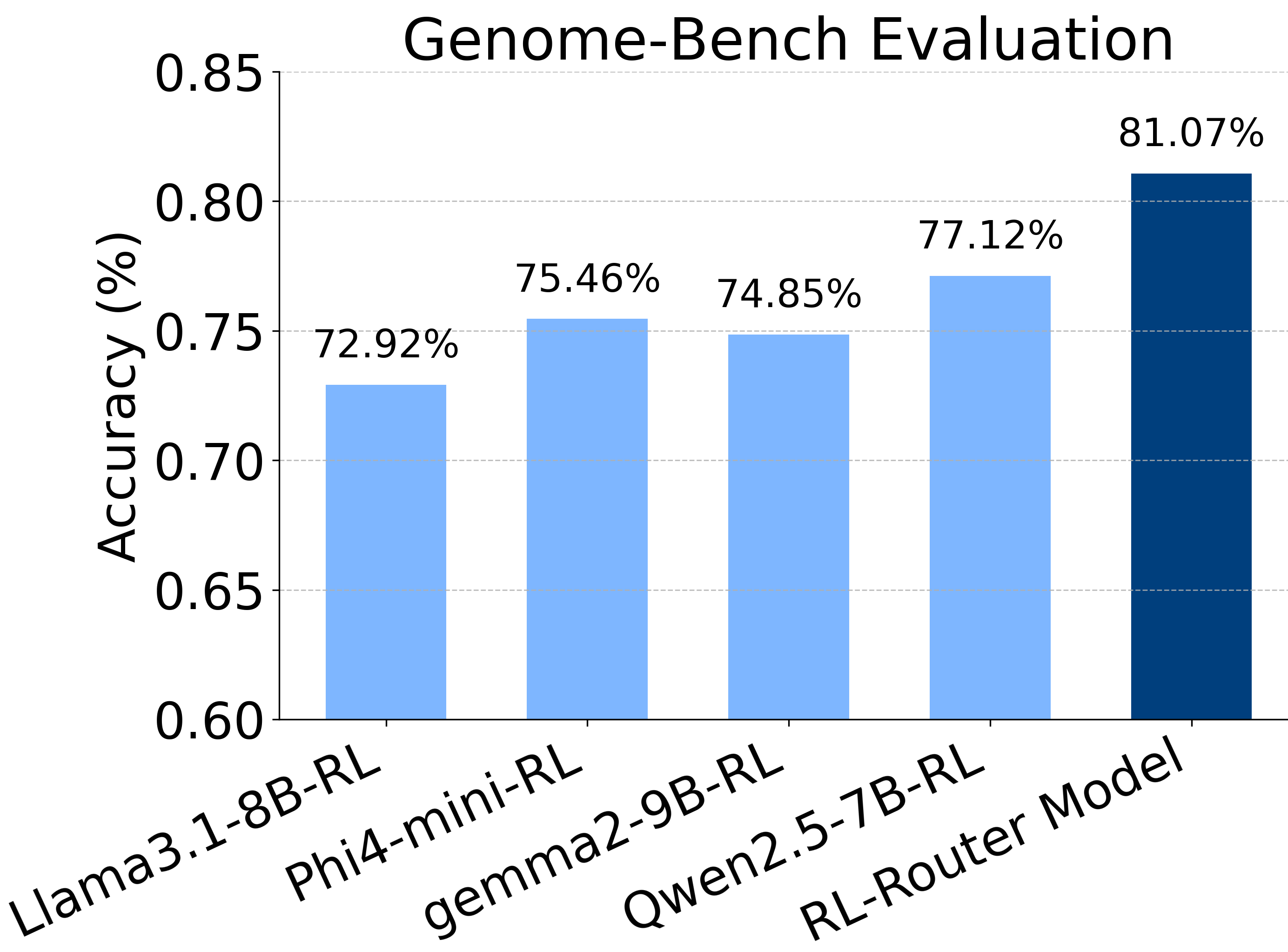

### router_performance_no_7B.png

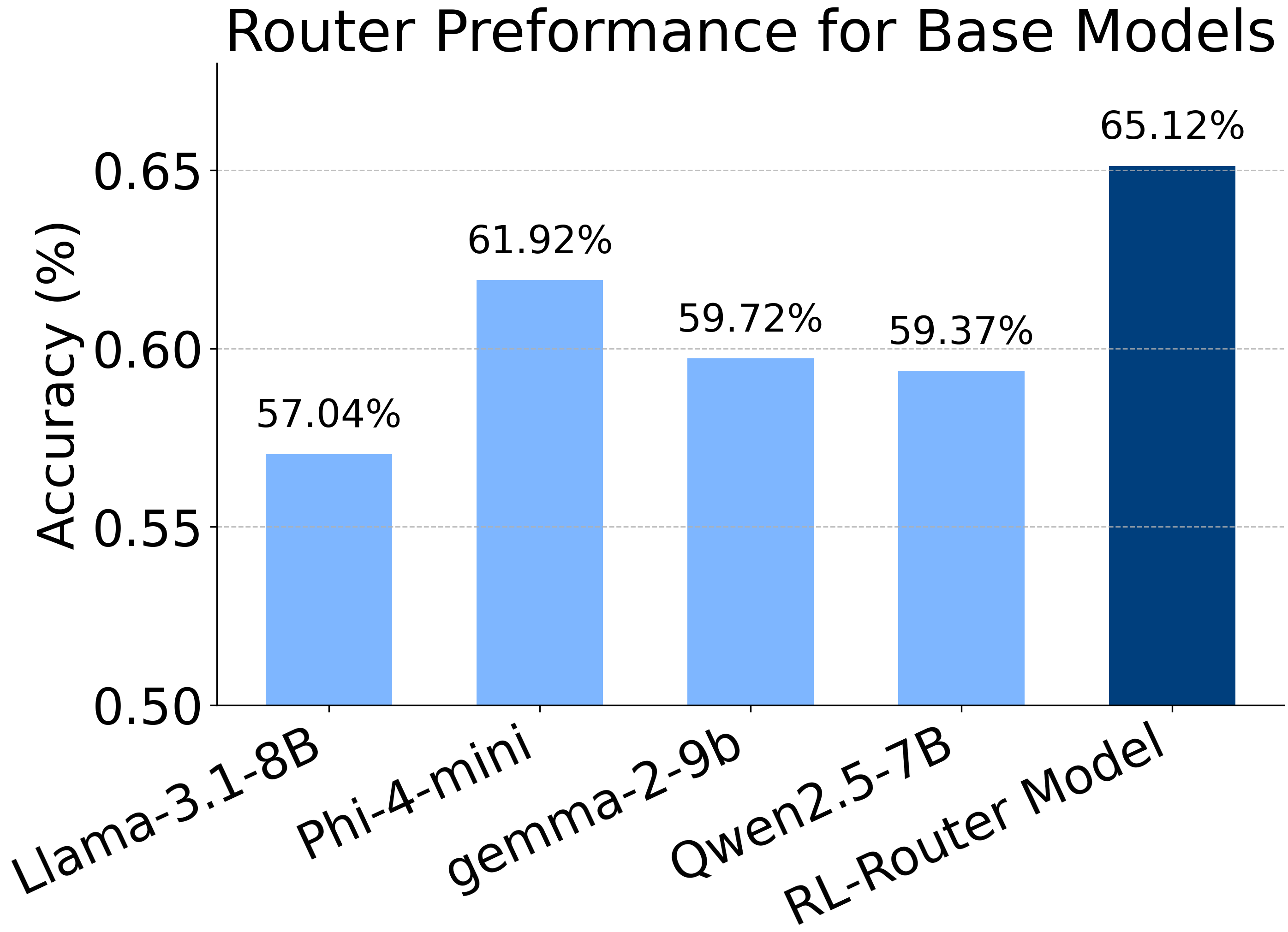

### temp1.pdf

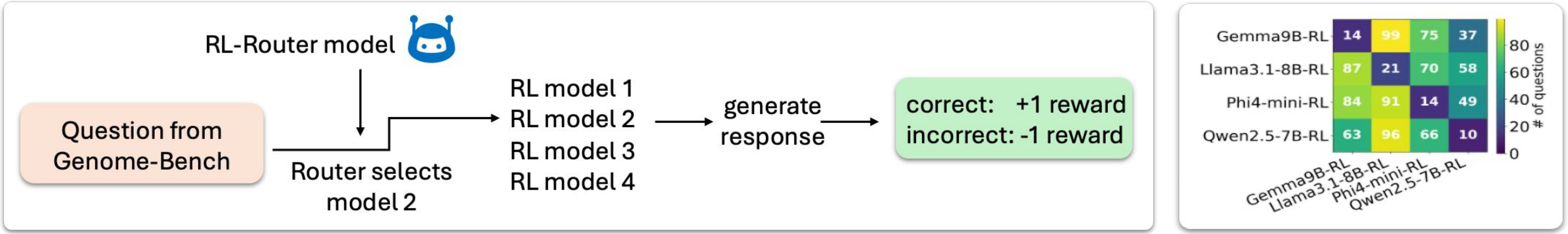

### temp.pdf

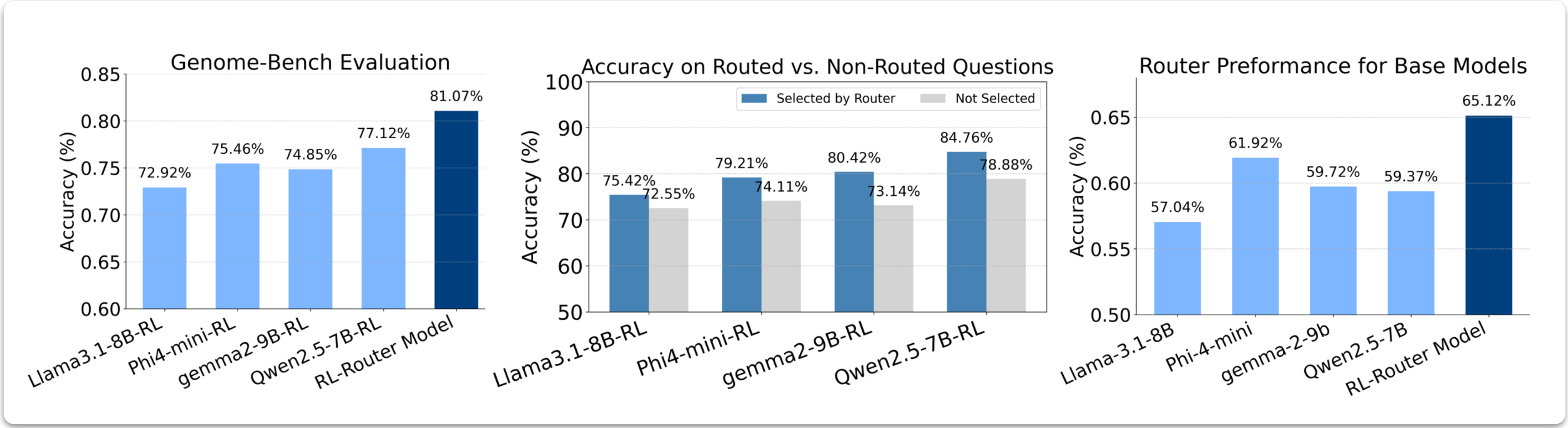
